# A mechanistic investigation of enhanced nitrogen use efficiency in wheat seedlings after treatment with an *Ascophyllum nodosum* biostimulant

**DOI:** 10.1101/2021.12.10.472083

**Authors:** Łukasz Łangowski, Oscar Goñi, Elomofe Ikuyinminu, Ewan Feeney, Shane O’Connell

## Abstract

Reduction in the emissions of the greenhouse gas nitrous oxide and nitrogen (N) pollution of ground water by improving nitrogen use efficiency (NUE) in crops is urgently required in pursuit of a sustainable agricultural future. Utilising an engineered biostimulant (PSI-362) derived from the brown seaweed Ascophyllum nodosum, we examined its effect on wheat seedling growth dynamics and mechanistic spatiotemporal changes at transcriptional and biochemical levels in relation to N uptake, assimilation and NUE. PSI-362-mediated biomass increase was associated with increased nitrate uptake and N assimilation in the form of glutamate, glutamine, free amino acids, soluble proteins and total chlorophyll. Phenotypical and biochemical analysis were supported by evaluation of differential expression of genetic markers involved in nitrate perception and transport (TaNRT1.1/NPF6.3), and assimilation (TaNR1 and TaNiR1, TaGDH2, TaGoGAT, TaGS1). Finally, a comparative analysis of the PSI-362 and two generic Ascophyllum nodosum extracts (ANEs) demonstrated that the NUE effect greatly differs depending on the ANE biostimulant used. In the current context of climate warming the transition of agriculture to a more sustainable model is urgently required. Application and adoption of precision biostimulants creates an opportunity for sustainable crop management, reduced production cost and environmental pollution, while maintaining yields.

## 1. Introduction

In order to maximise crop yields and economic return for the grower, large amounts of industrially produced N fertiliser are required every growing season in the form of nitrate (NO_3_-) and ammonium (NH_4_^+^) (Vance, 2001; Miller, AJ & Cramer, MD, 2005; Lassaletta, 2016; Yu & Zhuang, 2020). However, the production of N chemical fertilisers uses approximately 2% of global energy supply as fossil fuels, which contributes to greenhouse gas (GHG) emissions (Pfromm, 2017). Moreover, less than half of the added N fertiliser in the field is actually taken up by the crop (Raun & Johnson, 1999; White & Brown, 2010; Gutiérrez Rodrigo, 2012; Yan *et al.*, 2020). This low NUE increases the cost of crop production and has a negative environmental impact due to contamination of groundwater and GHG emissions by soil denitrification processes (Schaufler *et al.*, 2010; Chislock *et al.*, 2014; Kanter *et al.*, 2015; Wang *et al.*, 2019).

NUE encompasses the plant biological processes of N uptake, transport, assimilation and recycling from senescing organs, where each step is under strict regulation of environmental and genetic factors (Hirel & Krapp, 2020). Nitrate is the principal form of N taken up by plants and its availability in soil can fluctuate up to four orders of magnitude, either inhibiting or promoting root growth (Miller, A & Cramer, M, 2005; Bellegarde *et al.*, 2017; Liu *et al.*, 2017; Vidal *et al.*, 2020). Nitrate can also work as a signalling molecule and determines root architecture depending on the N status through modulating hormone homeostasis (Bouguyon *et al.*, 2016; Ristova *et al.*, 2016; Guan, 2017; Fu *et al.*, 2020). The perception of N occurs via the dual affinity transceptor NRT1.1, a nitrate sensor that is also functioning as a transporter in low and high N abundance (Fang *et al.*, 2021). From the soil, nitrate is predominantly taken up via NRT1/NPF and NRT2 families of transporters, which are known to have low and high affinity to this N form, respectively (Kiba & Krapp, 2016; Fang *et al.*, 2021). Inside the root and leaf cells nitrate undergoes reduction to nitrite by nitrate reductase (NR; EC 1.7.1.1), and subsequently to ammonium by nitrite reductase (NiR; EC 1.7.1.4) (Maia & Moura, 2014; Chamizo-Ampudia *et al.*, 2017). The assimilation of ammonium to glutamate (Glu) and glutamine (Gln) occurs through the sequential action of the enzymes glutamate dehydrogenase (GDH; EC 1.4.1.2), glutamine synthase (GS; EC 6.3.1.2) and glutamate synthase (NADH-GOGAT and Fd-GOGAT; EC 1.4.1.14 and 1.4.7.1, respectively) (Tercé-Laforgue *et al.*, 2013; Grzechowiak *et al.*, 2020). These N assimilation steps are also dependent and interlinked with central carbon (C) metabolism pathways such as the tricarboxylic acid cycle (TCA), that provides energy (ATP), reduced metabolic intermediates (NADPH) and carbon skeletons (2-oxoglutarate) (Masclaux-Daubresse *et al.*, 2010; Foyer & Noctor, 2011; Konishi *et al.*, 2014; Hirel & Krapp, 2020).

Increasing NUE can be achieved at multiple levels starting from improved soil management and fertiliser technologies. As the soil inorganic N pool is being depleted after every harvest, matching N supply with crop demand, in both space and time, is an effective way of improving NUE (Efretuei *et al.*, 2016). The use of novel fertilisers coated with bio-inhibitors and/or polymers is also a current alternative to reduce N losses when applied to soil (Snyder, 2017; Beig *et al.*, 2020; Dimkpa *et al.*, 2020; Gil-Ortiz *et al.*, 2020). Many transgenic approaches have also proved to be useful to enhance NUE through the overexpression of nitrate and ammonium transporters (e.g., NRT1/NPF, NRT2, and AMT1), the manipulation of *GS1* gene expression and altering the expression of transcription factors such as *NAC2* or *NLP7* to induce a broader impact on multiple N transporters or assimilation genes (He *et al.*, 2015; Yu *et al.*, 2015; O’Brien *et al.*, 2016; Gaudinier *et al.*, 2018; Sandhu *et al.*, 2021). However, most of these new varieties have not been validated yet in real field conditions and its implementation is still difficult from a regulation perspective and controversial in many countries (Wang, Q *et al.*, 2018; Wang, Y-Y *et al.*, 2018).

In recent years, the interest in plant biostimulants as sustainable solutions for crop production has grown significantly due to the development of next generation biostimulants with defined mode of action (MOA), backed by comprehensive research (du Jardin, 2015; Rouphael & Colla, 2020). In 2020 the global biostimulant market was worth approximately $2.2 billion and it is estimated that it will exceed $6.6 billion in 2031 (TransparencyMarketResearch, 2021). Specifically, plant biostimulants extracted from the brown seaweed *Ascophyllum nodosum* have a growing market share because of their ability to enhance tolerance to abiotic stress (Carmody *et al.*, 2020; Goñi *et al.*, 2021), improve crop quality traits (Frioni *et al.*, 2018; Łangowski *et al.*, 2019; Łangowski *et al.*, 2021), and enhance nutrient use efficiency in diverse crops (Jannin *et al.*, 2013; Billard *et al.*, 2014; Stamatiadis *et al.*, 2015; Laurent *et al.*, 2020; Goñi *et al.*, 2021). It is important to highlight that the biostimulant activity and composition of commercial *Ascophyllum nodosum* extracts (ANEs) is determined by the processing conditions of the raw material (Goñi *et al.*, 2018; Shukla *et al.*, 2019; Carmody *et al.*, 2020). However, there is still limited information explaining the in-depth mechanism of how ANEs can promote more N uptake and assimilation and the influence of processing parameters of the raw material with respect to the targeted enhancement of NUE in agronomic crops.

Recently, we examined the ability of an engineered biostimulant derived from *Ascophyllum nodosum* (PSI-362) to increase NUE in barley under field conditions. The targeted application of PSI-362 as a coating on a granular N mineral fertiliser allows up to 27% reduction in N fertiliser usage while maintaining or increasing crop yield through a defined physiological MOA (Goñi *et al.*, 2021). In the current study we focused on investigating the MOA of PSI-362 in winter wheat, a crop that provides more than 30% calories of global population (Li *et al.*, 2021). Wheat is characterized by lower maximum N recovery than barley (35-45%) and a requirement for a high level of N fertilisation to maintain rapid growth at early stages (Liao *et al.*, 2004; Zörb *et al.*, 2018; Sandhu *et al.*, 2021). Therefore, winter wheat seedlings (cv. Graham) tested in a controlled environment proved to be a reproducible plant system to assess the ability of PSI-362 biostimulant to enhance NUE using different phenotypic, biochemical, and molecular markers. The impact of PSI-362 regarding its dose, application time and its effect on different N supplementation rates on biomass growth dynamics, nitrate uptake, N assimilation products and differential expression of NUE related genes was evaluated. A comparative performance analysis between PSI-362 and two other commercial ANE biostimulants to improve NUE were also performed. The study demonstrates the efficacy of PSI-362 biostimulant in enhancing NUE in winter wheat through a defined MOA and demonstrates the benefits of next generation biostimulants that are precision engineered to resolve specific agronomic problems.

## 2. Material and Methods

### 2.1. Materials

Three commercially available liquid seaweed extracts of *A. nodosum* (PSI-362, ANE X and ANE Y) manufactured using different methods were applied to wheat plants as biostimulant treatments. ANE biostimulant containing the PSI-362 biomolecule complex is produced by Brandon Bioscience using a proprietary extraction at high temperatures and alkaline conditions (Goñi *et al.*, 2021). ANE X is manufactured using a proprietary process at low temperatures. ANE Y is manufactured using a proprietary process at high temperatures and alkaline pH. All chemical reagents used for the chemical characterization and biochemical assays were purchased from Sigma-Aldrich (Arklow, Ireland) and Bio-Rad (Watford, UK). The primers were purchased from Eurofins Genomics (Ebersberg, Germany).

### 2.2. Chemical characterization of ANE biostimulant treatments

Total N was determined according to the Dumas method. Total P and K was determined by acid digestion with nitric acid and quantification by ICP-MS. Ash, uronic acids, fucose, laminarin, free mannitol, and polyphenol content was evaluated according to (Goñi *et al.*, 2018). The content of other organic components was calculated by difference to the total organic amount.

### 2.3. Plant material, growth conditions, and ANE treatment application

The experiments involving wheat seedlings (cv. Graham) were performed in a growth chamber under controlled conditions (19/14°C with 16 hours of light and 8 hours of darkness and 80±5% RH under a light intensity of 120 μmol·m ^-2^os^-1^). Winter wheat seeds where sterilised with 0.5% v/v sodium hypochlorite for 5 min and subsequently rinsed 3 times with sterile water. Initially winter seeds were sown on 2% w/v agar medium without any supplementation for 6 days. Subsequently winter wheat seedlings were transferred in sterile conditions to transparent plastic containers (7 cm in diameter and 500 mL capacity) and grown over different durations (from 1 hour to 14 days) with and without treatments, unless specified otherwise. Depending on the experimental conditions, containers were filled with 150 ml of 2% agar w/v 1 MS, 1/10 MS or 0 MS medium. In order to test the potential impact of K present in PSI-362 on winter wheat seedlings development, two additional 2% w/v agar 0 MS medium controls were included, adding either 9.54 mg KCl or 11.14 mg K_2_SO_4_. For convenience and consistent treatment application, PSI-362, ANE X, and ANE Y were dried to powder at 105°C and mixed homogenously with liquid medium before agar gelling. As standard treatment conditions, 50 mg were added to 150 mL of liquid medium (0.33 mg·mL^-1^). For the concentration response experiments, 5 and 10 mg of PSI-362 were also added (0.03 and 0.06 mg·mL^-1^, respectively). For simplicity, the above mentioned concentrations are referred in the manuscript as 5 mg, 10 mg, and 50 mg. Due to space limitation in the plastic containers, the time course experiments at 1 hour, 1 day, 3 days, 6 days, and 9 days of treatment were performed using half the volume of medium (75 mL) while maintaining the same concentration of PSI-362 dose (25 mg added to 75 mL). Experiments growing plants for 6 and 14 days on 0 MS were also performed in 75 mL medium. At the end of the experiment, wheat seedlings were harvested and total fresh weight (FW) was determined. Shoots and roots of the seedlings were separated to determine their FW. Seedling samples were snap-frozen in liquid nitrogen, ground and kept in −80°C until further biochemical and gene expression analysis. The efficiency of the conversion of added inorganic N into total plant FW was calculated for control and treated plants in each plastic container as below:

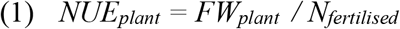

### 2.4. RNA extraction and RT-qPCR

Total RNA was isolated from frozen ground wheat seedlings using a Total RNA Purification Kit (Norgen Biotek, Canada) as per the manufacturer’s instructions. RNA was treated with RNase-Free DNase I Kit (Norgen Biotek, Canada) in order to efficiently remove genomic DNA contamination. RNA concentration and purity was measured using Qubit (Thermo Fisher Scientific). Expression analysis of genes *TaNRT1.1/NPF6.3* (Buchner *et al.*, 2015), *TaNR1* (Buchner *et al.*, 2015), *TaNiR1* (Buchner *et al.*, 2015), *TaGDH2* (Buchner *et al.*, 2015), *TaGS1* (Buchner *et al.*, 2015), *TaNADH-GoGAT* (Li *et al.*, 2020) and *TaAMT1c* (Li *et al.*, 2017) was performed by RT-qPCR using a Roche LightCycler® 96 System (Roche, UK) and a LightCycler® RNA Master SYBR Green I one-step kit (Roche, UK) according to the manufacturer’s instructions. The relative expression level of the above mentioned genes was calculated relative to reference genes *TaACT2* (Bajgain *et al.*, 2018), *TaBeta-Tubulin* (Bajgain *et al.*, 2018), and *TaEF1a* (Li *et al.*, 2017) from which the most stable to PSI-362 treatment was *TaEF1a.* The 2^-ΔΔCT^ method was used to quantify relative normalized gene expression levels. The primers sequences used are shown in Table S1.

### 2.5. Measurement of N metabolites, soluble protein, and photosynthetic pigments

Nitrate content was determined in material grinded from entire wheat seedlings according to the protocol described by (Goñi *et al.*, 2021). Nitrate content was also evaluated in 1/10 MS medium at the end of the experiment. To extract the nitrate from the medium 100 mg sample of gel from the bottom of the container was placed in a plastic eppendorf tube, grinded with a plastic pistil and subsequently incubated for 15min at 50°C in order to completely dissolve the gel. Once a liquid sample was obtained, an aliquot was used to measure nitrate content using the spectrophotometric method described above (Goñi *et al.*, 2021). Total FAA and ammonium extraction was performed by mixing 30 mg of grinded tissue with 700 μl of ethanol 70% (v/v) according to (Goñi *et al.*, 2021). These supernatants were first used for the estimation of total free amino acids content using ninhydrin reagent and a calibration standard curve with glutamic acid (Goñi *et al.*, 2021). Glutamic acid, glutamine, and ammonium content was evaluated in the same extracts after derivatization with DEEMM through RP-HPLC and UV detection at 280 and 300 nm (Gómez-Alonso *et al.*, 2007). Soluble protein and photosynthetic pigments from entire wheat seedlings were measured according to (Goñi *et al.*, 2021). All biochemical markers were expressed with respect to sample FW.

### 2.6. Statistical analysis

Chemical analysis of commercial ANEs was performed on a minimum number of 3 biological replicates, using 3 technical replicates per biological replicate. Phenotypic and NUE assessment of the winter wheat grown in plastic containers was performed using 9 independent biological replicates, with 8 plants per replicate. Assessment of N metabolites, soluble protein, and photosynthetic pigments was performed in at least 4 independent biological replicates, using 3 technical replicates per biological replicate. For gene expression analysis, at least 3 biological replicates of each sample was performed, using 3 technical replicates per biological replicate.

Statistics were evaluated with Sigma Plot 12 and Statgraphics Centurion XVI software. The effect on one single factor was analysed using one-way analysis of variance (ANOVA) by Student–Newman–Keuls’ method at *p* ≤ 0.05. The effect of two factors was determined using the two-way ANOVA by Student–Newman–Keuls’ method at *p* ≤ 0.05. Differences between respective control and PSI-362 treatment were analysed using t-test at *p* ≤ 0.05 for the interaction between two factors. The statistical effect of each factor was also evaluated separately, comparing the respective means through t-test or one-way ANOVA Student– Newman–Keuls’ method at *p* ≤ 0.05. The application of all parametric tests was performed after checking the normality of the data (Shapiro-Wilk’s test) and equal variance assumptions. Unless stated otherwise, all data are expressed as average ± standard error (SE). Details of the individual sample size for each analysis and statistical test used is mentioned in the tables and figure legends.

## 3. RESULTS

### 3.1. PSI-362 enhances NUE in winter wheat in concentration dependent manner

In order to test the efficacy of PSI-362 in enhancing NUE in winter wheat, 6 days old seedlings were transferred to agar medium supplemented with reduced N medium (1/10 MS) and three doses of PSI-362 in powder form (5 mg, 10 mg, and 50 mg, respectively). After 6 days of treatment, total plant, shoot and root biomass, and nitrate content in fresh tissue was measured (Fig. 1). All treated plants showed a significant increase in all biomass and nitrate content parameters with respect to the control. A clear dose response to PSI-362 was evident. Total plant biomass increases of 10.7%, 17.7%, and 66% were recorded when PSI-362 rate was augmented from 5 mg to 10 mg, and finally 50 mg (*p* = 0.08, 0.024, and ≤ 0.001, respectively) (Fig. 1c, Table 1). These variations were also reflected in enhanced NUE values of treated plants compared to control (Table S2). The application of PSI-362 at 3 different rates had a stronger positive effect on root biomass than on shoot tissue. While the application of 5 mg, 10 mg and 50 mg concentrations stimulated shoot biomass by 5.7%, 11.7%, and 33.6% compared to untreated control (*p* = 0.194, 0.046, and ≤ 0.001 respectively), the same treatments increased root biomass by 18.6%, 36.3%, and 142.2% (*p* = 0.092, 0.014, and ≤ 0.001 respectively) (Fig. 1c, Table 1). As for the nitrate uptake, the lowest concentration of PSI-362 was effective in increasing this metabolite level by 31.7% compared to control (*p* = 0.001). This effect was even higher with increased rates of PSI-362 (10 mg and 50 mg), which increased plant nitrate content by 48.9% and 89.9%, respectively (*p* ≤ 0.001) (Fig.1d, Table 1).

**Fig. 1.**
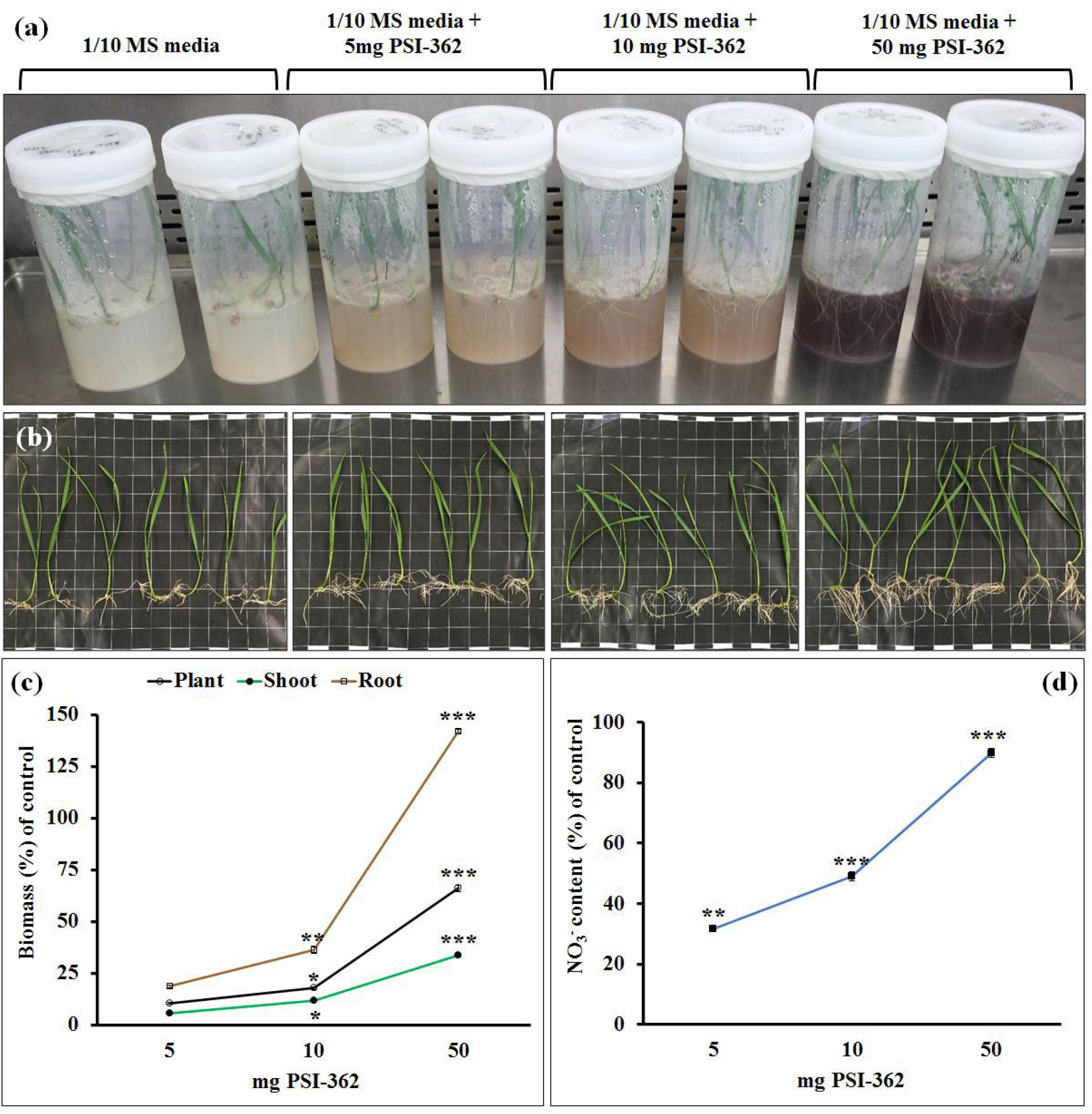
PSI-362 enhances plant biomass and nitrate uptake in winter wheat in concentration dependent manner. The image presents an example of the experimental setup used in this study. Plastic containers with lids sealed with Micropore^TM^ tape contain 12-day-old winter wheat seedlings (cv. Graham) treated for 6 days with 0 mg (control), 5 mg, 10 mg, and 50 mg of PSI-362 in 150ml 1/10 MS agar medium (a). Pictures in the lower row represent 12-day-old winter wheat seedlings (cv. Graham) grown in plastic containers according to conditions mentioned above (b). Charts represent total plant, shoot, and root biomass change (c) and nitrate content variation (d) with respect to the control. Each treatment was performed using 9 independent biological replicates, with 8 plants per replicate. Nitrate content assessment was performed with at least 4 independent biological replicates. Means followed by asterisk (***, **, and * significant at *p* ≤ 0.001, *p* ≤ 0.01, and *p* ≤ 0.05, respectively) indicate statistically significant differences for each parameter between control and PSI-362 treatment.

**Table 1.**
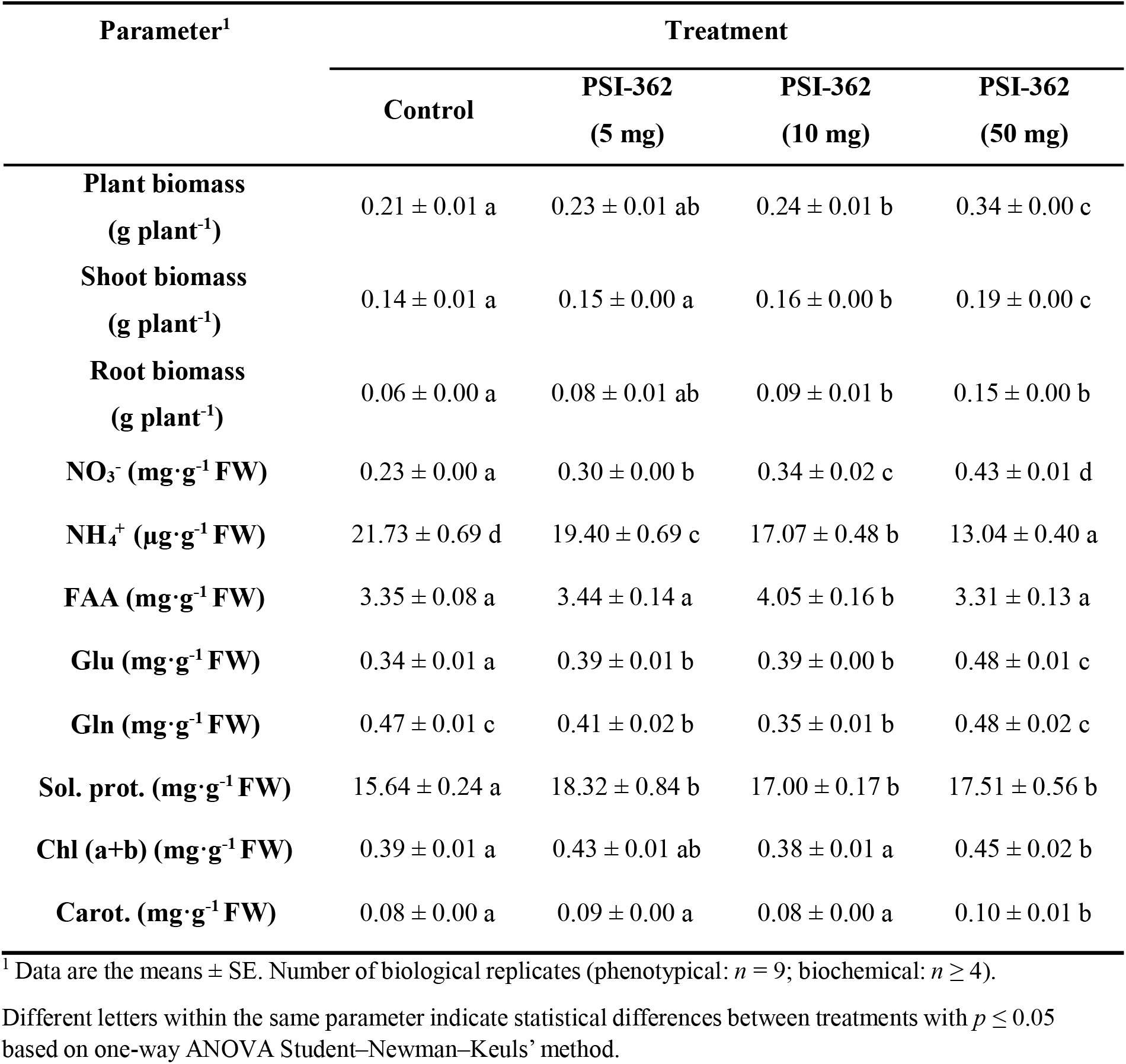
Effect of PSI-362 concentration applied for 6 days on phenotypic and biochemical parameters in winter wheat seedlings (cv. Graham) grown on reduced N media (1/10 MS).

|  | Control | PSI-362<br>(5 mg) | PSI-362<br>(10 mg) | PSI-362<br>(50 mg) |
| --- | --- | --- | --- | --- |
| **Plant biomass**<br>(g plant<sup>-1</sup>) | 0.21 ± 0.01 a | 0.23 ± 0.01 ab | 0.24 ± 0.01 b | 0.34 ± 0.00 c |
| **Shoot biomass**<br>(g plant<sup>-1</sup>) | 0.14 ± 0.01 a | 0.15 ± 0.00 a | 0.16 ± 0.00 b | 0.19 ± 0.00 c |
| **Root biomass**<br>(g plant<sup>-1</sup>) | 0.06 ± 0.00 a | 0.08 ± 0.01 ab | 0.09 ± 0.01 b | 0.15 ± 0.00 b |
| **NO<sub>3</sub><sup>-</sup> (mg·g<sup>-1</sup> FW)** | 0.23 ± 0.00 a | 0.30 ± 0.00 b | 0.34 ± 0.02 c | 0.43 ± 0.01 d |
| **NH<sub>4</sub><sup>+</sup> (μg·g<sup>-1</sup> FW)** | 21.73 ± 0.69 d | 19.40 ± 0.69 c | 17.07 ± 0.48 b | 13.04 ± 0.40 a |
| **FAA (mg·g<sup>-1</sup> FW)** | 3.35 ± 0.08 a | 3.44 ± 0.14 a | 4.05 ± 0.16 b | 3.31 ± 0.13 a |
| **Glu (mg·g<sup>-1</sup> FW)** | 0.34 ± 0.01 a | 0.39 ± 0.01 b | 0.39 ± 0.00 b | 0.48 ± 0.01 c |
| **Gln (mg·g<sup>-1</sup> FW)** | 0.47 ± 0.01 c | 0.41 ± 0.02 b | 0.35 ± 0.01 b | 0.48 ± 0.02 c |
| **Sol. prot. (mg·g<sup>-1</sup> FW)** | 15.64 ± 0.24 a | 18.32 ± 0.84 b | 17.00 ± 0.17 b | 17.51 ± 0.56 b |
| **Chl (a+b) (mg·g<sup>-1</sup> FW)** | 0.39 ± 0.01 a | 0.43 ± 0.01 ab | 0.38 ± 0.01 a | 0.45 ± 0.02 b |
| **Carot. (mg·g<sup>-1</sup> FW)** | 0.08 ± 0.00 a | 0.09 ± 0.00 a | 0.08 ± 0.00 a | 0.10 ± 0.01 b |

<sup>1</sup> Data are the means ± SE. Number of biological replicates (phenotypical: $n = 9$ ; biochemical: $n \geq 4$ ).
Different letters within the same parameter indicate statistical differences between treatments with $p \leq 0.05$ based on one-way ANOVA Student–Newman–Keuls' method.

### 3.2. PSI-362 efficacy under diverse N conditions

To determine the effect of distinct N supplementation conditions on PSI-362 performance in winter wheat seedlings, high N (1 MS) and N deficient (1/10 MS) media were tested (Fig. 2). The two-way ANOVA showed that PSI-362 treatment, N rate and the interaction between factors (PxN) were statistically significant for plant, shoot and root biomass (Table 2). As observed previously, PSI-362 treated plants grown for 6 days in reduced N medium (1/10 MS) showed remarkable total plant, shoot and root biomass increase with respect to control (63.9%, 32.1%, and 132.6%, respectively; *p* ≤ 0.001). It is important to note that plants treated with 50 mg of PSI-362 and supplemented with high N levels (1 MS) also presented a statistically significant increase in total plant, shoot and root biomass increase with respect to the relevant control (29.9%, 18.6%, 49.3%; *p* ≤ 0.001) (Fig. 2e). Interestingly, PSI-362 treated plants grown in reduced N level (1/10 MS) also outperformed control plants grown on medium with 10-fold higher N content (1 MS), closing a 90% N gap at phenotypic level. PSI-362 treated plants showed similar total plant, shoot and root biomass absolute values under both N regimes (Table 2) indicating that wheat seedlings, even when grown under high N level, cannot exceed certain plant growth dynamics and NUE improvements due to genetic limitations (Table S3).

**Fig. 2.**
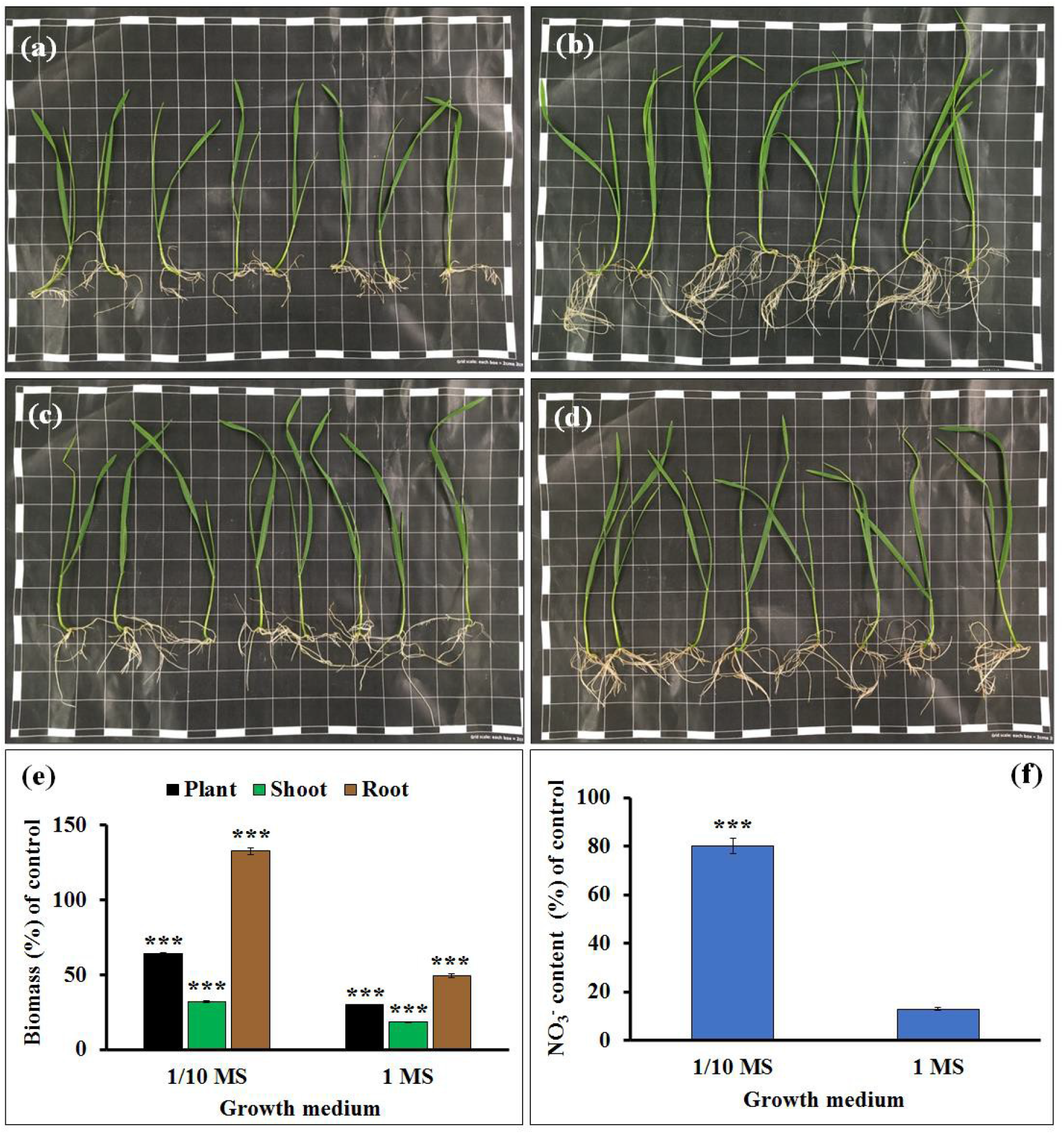
PSI-362 efficacy on plant biomass and nitrate uptake under diverse N conditions. Pictures represent 12-day-old control (a, c) and 50 mg PSI-362 treated (b, d) winter wheat seedlings (cv. Graham) grown at different N supplementation, 1/10 MS (a, b), and 1 MS agar medium (c, d). Charts represent total plant, shoot, and root biomass change (e) and nitrate content variation (f) with respect to the control. Each treatment was performed using 9 independent biological replicates, with 8 plants per replicate. Nitrate content assessment was performed with at least 4 independent biological replicates. Since interaction PxN was significant (* *p* ≤ 0.05), the data was subjected to t-test, comparing PSI-362 treatment versus control within the same N rate. In this case, means followed by asterisk (*** significant at *p* ≤ 0.001) indicate statistically significant differences between control and PSI-362 treatment for each parameter within the same N rate.

**Table 2.**
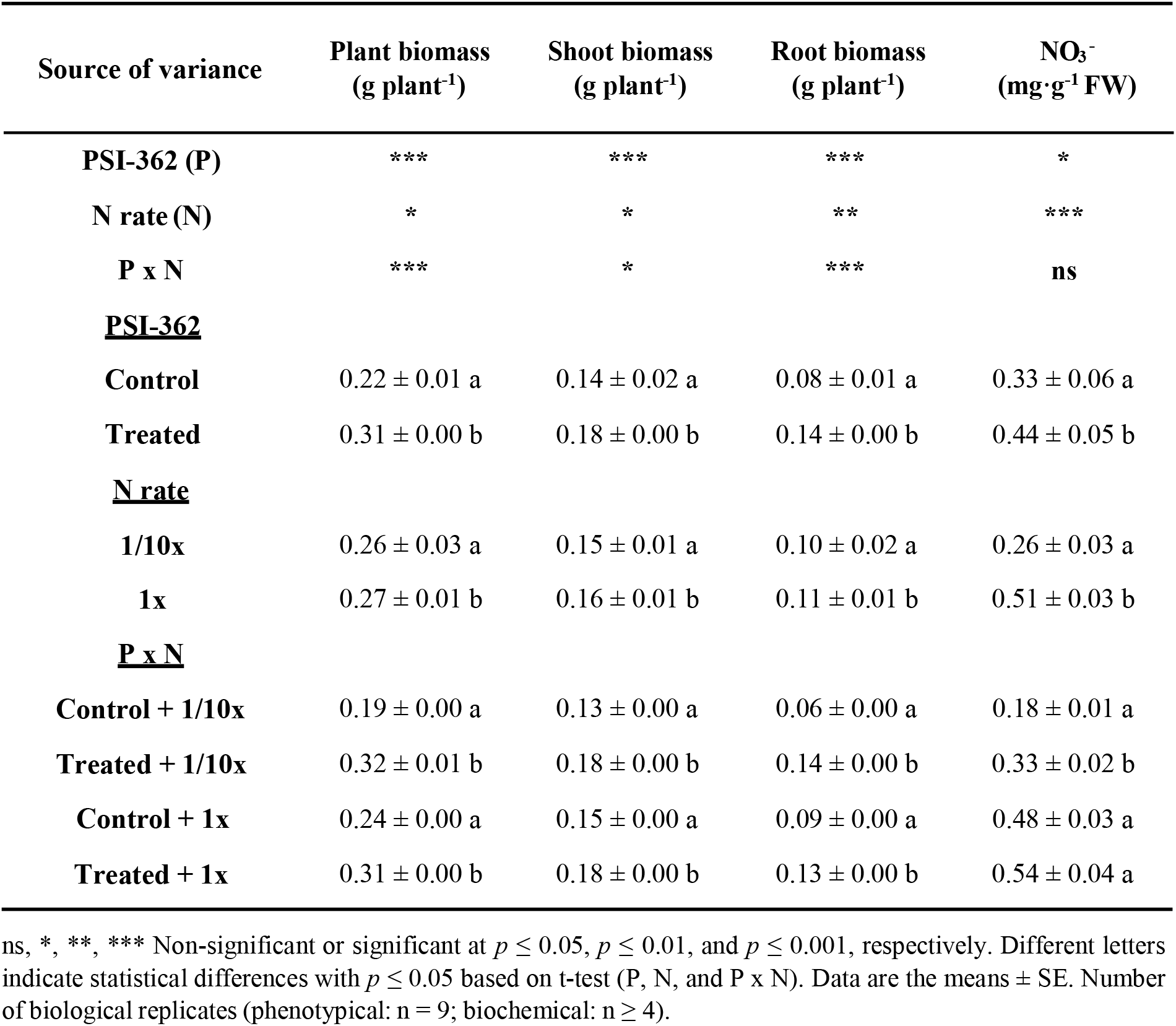
Analysis of variance and mean comparisons for phenotypic parameters and nitrate content in winter wheat seedlings (cv. Graham) treated with PSI-362 for 6 days and growing under two different N rates

In terms of nitrate content in fresh tissue, the recorded concentration was positively dependent on N availability (*p* ≤ 0.001) and PSI-362 treatment (*p* = 0.012). The lowest concentration detected was in control plants grown on 1/10 MS, while the highest one was in plants grown on 1 MS medium supplemented with PSI-362 (Table 2). Interestingly, there was no statistically significant interaction between factors (P x N) in nitrate content, indicating that PSI-362 had a general stimulating effect by increasing N uptake by 80% (*p* ≤ 0.001) and 12.9% (*p* = 0.217) in plants growing under reduced (1/10 MS) and full (1 MS) N rate, respectively (Fig. 2e, Table 2). This experiment shows that, although there is a clear correlation between enhanced nitrate uptake and increased biomass in treated plants with PSI-362, absolute nitrate content values does not necessarily correlate with total plant biomass when plants are grown at different N levels.

### 3.3. PSI-362 NUE effect is rapid and long-lasting

In order to test how quickly wheat seedlings respond to PSI-362 treatment and what the temporal dynamics of the growth is, biomass and nitrate parameters were recorded after 1, 3, 6, and 9 days of stimulation with 50 mg PSI-362 in reduced N medium (1/10 MS) (Table 3, Fig. S1). After 24h of treatment, PSI-362 did not induce any statistically significant change in plant growth versus the control, with a slight decrease in plant and shoot biomass (−3.6% and −7.5%; *p* = 0.713 and 0.525, respectively) and a small increase in root biomass (+4.1%, *p* = 0.824) (Fig. 3a). In addition, there was a statistically significant interaction between factors (P x T) in nitrate content, being characterized by a moderate increase in treated plants after 1 day (+12.2%; *p* ≤ 0.001). This suggests that PSI-362 initially increases the nitrate uptake capacity through existing channels (Fig. 3b). Three, six, and nine days of treatment with PSI-362 led to gradual total plant biomass and NUE increase in comparison to respective control (21.7%, 44.3%, and 39.3, respectively) (*p* ≤ 0.001) (Fig. 3a, Table S4). Shoot and root biomass measured after each time point demonstrated that the boosted shoot growth rate with respect to control was plateauing around 20% after 9 days of treatment while root growth rate was still increasing by 92% (Fig. 3a, Fig. S1). The nitrate content analysis of fresh tissue revealed that the highest relative nitrate level between treated and control plants was recorded after 3 days (+71.3%), declining slightly after 6 days (+60.6%), and reaching a similar level as that observed after 1 day of treatment with PSI-362 (+15.0%) in plants treated for 9 days (Fig. 3b).

**Table 3.** Analysis of variance and mean comparisons for phenotypic parameters and nitrate content in winter wheat seedlings (cv. Graham) grown on reduced N media (1/10 MS) and treated with PSI-362 during different time points.

| Source of variance | Plant biomass<br>(g plant<sup>-1</sup>) | Shoot biomass<br>(g plant<sup>-1</sup>) | Root biomass<br>(g plant<sup>-1</sup>) | NO<sub>3</sub><sup>-</sup><br>(mg·g<sup>-1</sup> FW) |
| --- | --- | --- | --- | --- |
| **PSI-362 (P)** | *** | *** | *** | *** |
| **Time (T)** | *** | *** | *** | *** |
| **P x T** | *** | ** | *** | *** |
| **<u>PSI-362</u>** |  |  |  |  |
| **Control** | 0.16 ± 0.01 a | 0.11 ± 0.01 a | 0.05 ± 0.00 a | 0.20 ± 0.01 a |
| **Treated** | 0.22 ± 0.02 b | 0.13 ± 0.01 b | 0.09 ± 0.01 b | 0.31 ± 0.02 b |
| **<u>Time</u>** |  |  |  |  |
| **1 d** | 0.09 ± 0.00 a | 0.06 ± 0.00 a | 0.03 ± 0.00 a | 0.14 ± 0.00 a |
| **3 d** | 0.17 ± 0.01 b | 0.11 ± 0.00 b | 0.06 ± 0.01 b | 0.24 ± 0.02 b |
| **6 d** | 0.24 ± 0.02 c | 0.14 ± 0.01 c | 0.10 ± 0.01 c | 0.35 ± 0.03 c |
| **9 d** | 0.09 ± 0.00 d | 0.18 ± 0.01 d | 0.08 ± 0.01 d | 0.29 ± 0.01 b |
| **<u>P x T</u>** |  |  |  |  |
| **Control + 1 d** | 0.09 ± 0.00 a | 0.06 ± 0.00 a | 0.03 ± 0.00 a | 0.13 ± 0.00 a |
| **Treated + 1 d** | 0.09 ± 0.00 a | 0.06 ± 0.00 a | 0.03 ± 0.00 a | 0.14 ± 0.00 b |
| **Control + 3 d** | 0.15 ± 0.00 a | 0.11 ± 0.00 a | 0.05 ± 0.00 a | 0.18 ± 0.00 a |
| **Treated + 3 d** | 0.19 ± 0.00 b | 0.11 ± 0.01 a | 0.08 ± 0.00 b | 0.30 ± 0.01 b |
| **Control + 6 d** | 0.19 ± 0.00 a | 0.13 ± 0.00 a | 0.07 ± 0.00 a | 0.27 ± 0.01 a |
| **Treated + 6 d** | 0.28 ± 0.01 b | 0.15 ± 0.00 b | 0.12 ± 0.00 b | 0.43 ± 0.04 b |
| **Control + 9 d** | 0.22 ± 0.00 a | 0.16 ± 0.01 a | 0.06 ± 0.01 a | 0.30 ± 0.01 a |
| **Treated + 9 d** | 0.31 ± 0.00 b | 0.20 ± 0.01 b | 0.11 ± 0.00 b | 0.35 ± 0.01 b |

\*\*, \*\*\* Significant at $p \leq 0.01$ and $p \leq 0.001$ , respectively. Different letters indicate statistical differences with $p \leq 0.05$ based on t-test (P and P x T) or Student–Newman–Keuls’ method (T). Data are the means ± SE. Number of biological replicates (phenotypical: n = 9; biochemical: n ≥ 4).

**Fig. 3.**
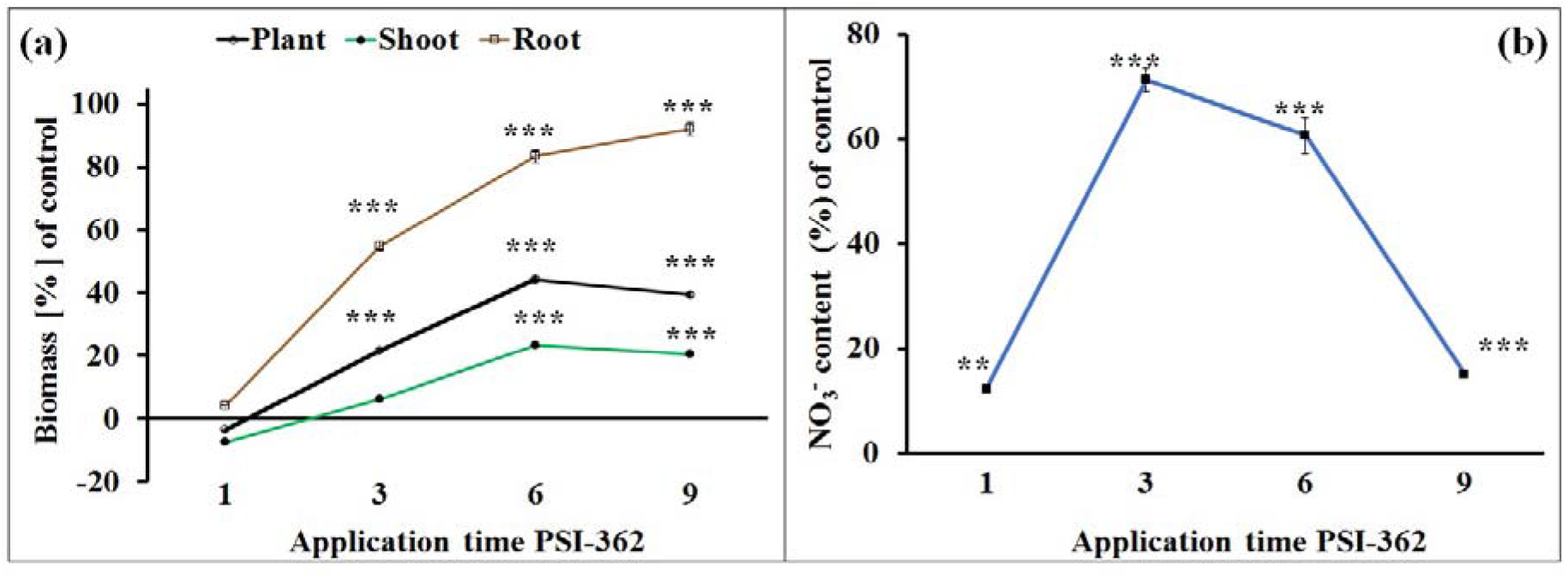
Effects of PSI-362 treatment on plant biomass and nitrate uptake during different time points. 6-day-old winter wheat seedlings treated with 0 mg (control) and 25 mg PSI-362 in 75ml 1/10 MS agar medium for 1, 3, 6, and 9 days. Charts represent total plant, shoot, and root biomass change (a) and nitrate content variation (b) with respect to the control. Each treatment was performed using 9 independent biological replicates, with 8 plants per replicate. Nitrate content assessment was performed with at least 4 independent biological replicates. Since interaction PxT was significant (* p ≤ 0.05), the data was subjected to one-way ANOVA Student-Newman-Keuls’ method, comparing PSI-362 treatment versus control within the same application time. In this case, means followed by asterisk (*** significant at p ≤ 0.001) indicate statistically significant differences between control and PSI-362 treatment for each parameter within the same application time.

Such temporal dynamics in biomass and nitrate uptake can be explained by either depletion of the nutrients in reduced N medium or the nature of the biostimulant treatment. To answer this question, the nitrate level was also measured in 1/10 MS agar media after collecting treated and untreated plants grown for 3, 6, and 9 days (Fig. 4). As expected, the nitrate content declined with each recorded day, however this nutrient declined more rapidly in growth media supplemented with PSI-362. The most significant nitrate drop compared to control was recorded after 6 days (−48.9%; *p* ≤ 0.001) (Fig. 4), coinciding with the highest relative biomass increase (+44.3%) (Fig. 3a). After 9 days of PSI-362 treatment, the nitrate intake dynamics in the growth medium between control and treated plants were still significantly different despite apparent nutrient depletion in treated plants (−31.2%; *p* ≤ 0.05) (Fig. 4).

**Fig. 4.**
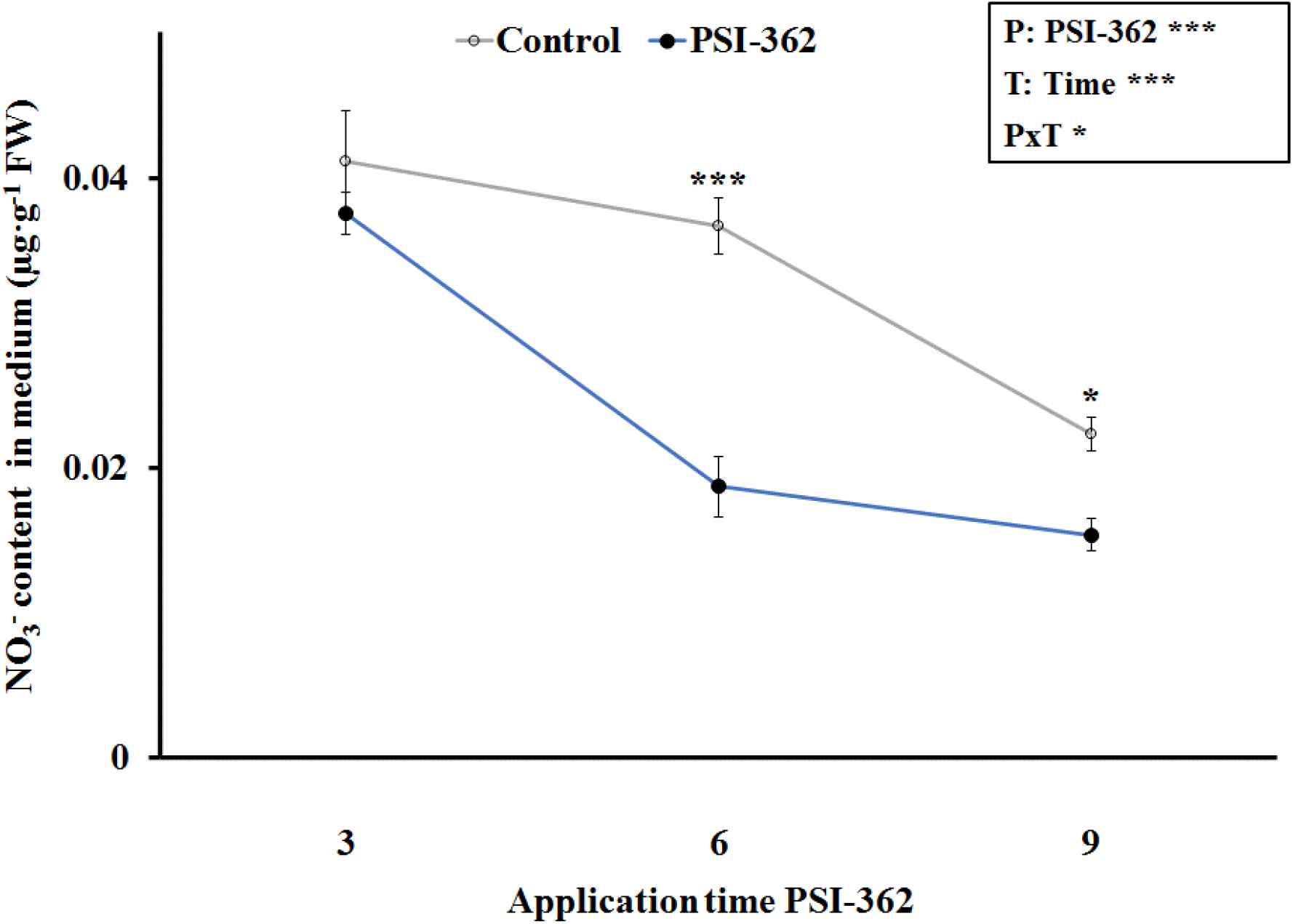
Effect of PSI-362 treatment on nitrate content in 1/10 MS agar medium during different time points. 6-day-old winter wheat seedlings were treated with 0 mg (control) and 25 mg PSI-362 in 75ml 1/10 MS agar medium for 3, 6, and 9 days. Nitrate content was assessed in control and PSI-362 supplemented 1/10 MS agar medium recovered from plastic containers using at least 4 independent biological replicates. Since interaction PxT was significant (* p ≤ 0.05), the data was subjected to t-test, comparing PSI-362 treatment versus control within the same application time. In this case, means followed by asterisk (*** and * significant at p ≤ 0.001 and p ≤ 0.05, respectively) indicate statistically significant differences between control and PSI-362 treatment within the same application time.

### 3.4. PSI-362 promotes N uptake from the grain and does not work like a nutrient

In order to gain more insight into the mode of action of PSI-362, 6-day-old winter wheat seedlings were transferred on 0 MS medium supplemented with and without PSI-362 for 6 and 14 days. The total plant biomass of the control and PSI362 plants at both time points was determined. After 6 days growing on 0 MS media, total plant biomass of the control and treated plants were lower in comparison to those previously grown on 1/10 MS for the same period of time (−9.3% and −25.1%, respectively) (Fig. S2, Table 3). If PSI-362 was only enhancing uptake of N provided in MS media, treated and untreated plants grown on 0 MS should show the same growth dynamic. However, treated plants after 6 and 14 days on 0 MS medium were significantly bigger than their respective controls, increasing their biomass by 19.0% and 38.5%, respectively; (*p* ≤ 0.001) (Fig. S2). It indicates that PSI-362 is not only improving NUE when N mineral nutrients are available in the medium, but most likely also improving the N uptake and transport from the germinated grain, that in this experimental setup, is the only N source.

In order to evaluate the possibility of whether PSI-362 ANE biostimulant can work as a source of nutrients, its macronutrient NPK composition was determined and compared to nutrients added with MS media. The overall NPK content in PSI-362 was 0.4:0.1:8.0% w/v, which provided 0.396 mg N, 0.099 mg P, and 8.267 mg K in 50 mg of PSI-362 supplemented to 150 mL of 1/10 MS medium (Table 4). The amount of total N present in 1/10 MS medium is 12.6 mg per 150 mL, therefore PSI-362 addition increased the N content by a marginal 3.2%, which is not sufficient to account for the observed gains in biomass or nitrate uptake. The fact that PSI-362 treated plants grown on 1/10 MS had similar biomass values to those grown on 10-fold more N rich medium, or 34.4% higher total plant biomass than control plants on 1 MS, also suggests that PSI-362 does not have a significant role as N nutrient (Table 2). Furthermore, treated plants grown on 0 MS for 6 days gained more total plant biomass than control plants grown on 1/10 MS (+17.4%; *p* ≤ 0.001) (Fig. 5a).

**Table 4.** Compositional analysis of ANE biostimulants

| Component <sup>1</sup> | Treatment |  |  |
| --- | --- | --- | --- |
|  | PSI-362 <sup>3</sup> | ANE X | ANE Y |
| **Total N % (w/v) <sup>2</sup>** | 0.41 ± 0.03 b | 0.20 ± 0.02 a | 0.21 ± 0.03 a |
| **Total P % (w/v)** | 0.10 ± 0.01 b | 0.01 ± 0.01 a | 0.02 ± 0.01 a |
| **Total K % (w/v)** | 8.03 ± 0.34 c | 0.59 ± 0.04 a | 1.02 ± 0.09 b |
| **Ash % (w/w) <sup>3</sup>** | 44.02 ± 0.33 b | 20.51 ± 0.23 a | 46.71 ± 0.30 c |
| **Uronic acids % (w/w)** | 7.16 ± 0.51 b | 5.68 ± 0.29 a | 11.54 ± 0.10 c |
| **Fucose % (w/w)** | 8.04 ± 0.05 c | 3.88 ± 0.15 a | 6.06 ± 0.05 b |
| **Laminarin % (w/w)** | 1.57 ± 0.04 a | 9.56 ± 0.22 b | 1.35 ± 0.02 a |
| **Free mannitol % (w/w)** | 4.13 ± 0.07 a | 6.20 ± 0.06 b | 4.00 ± 0.14 a |
| **Polyphenol % (w/w)** | 13.68 ± 0.18 c | 4.85 ± 0.07 a | 8.37 ± 0.14 b |
| **Other organic % (w/w)** | 21.40 ± 0.06 a | 49.32 ± 0.29 b | 21.97 ± 0.12 a |

<sup>1</sup> Data are the means ± SE. Number of biological replicates ( $n \geq 3$ ).
<sup>2</sup> ANEs NPK analysis is expressed with respect to their liquid content.
<sup>3</sup> ANEs chemical compositional analysis is expressed with respect to their dry weight.
Different letters within the same parameter indicate statistical differences between treatments with $p \leq 0.05$ based on one-way ANOVA Student-Newman-Keuls' method.

**Fig. 5.**
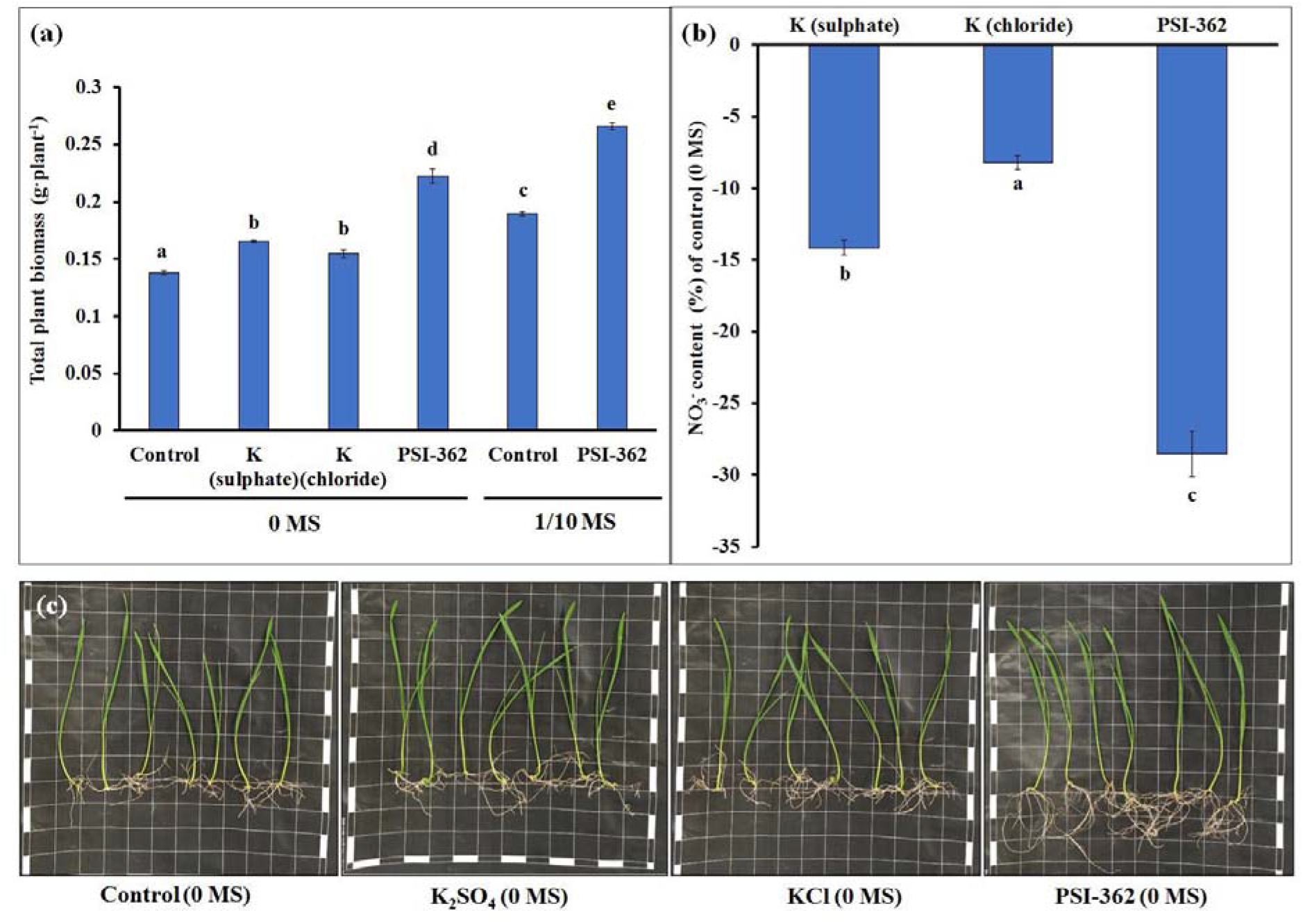
Effect of PSI-362 treatment in winter wheat seedlings grown in nutrient starving conditions. 6-day-old winter wheat seedlings were treated with 0 mg (control) and 50 mg PSI-362 in 150 ml 0 MS and 1/10 MS agar medium for 6 days. Additional 1/10 MS controls was included adding 14,64 mg of total K in the form of K2CO3. Charts represent total plant biomass of different treatments (a) and nitrate content variation with respect to 0 MS and 1/10 MS control (b). Each treatment was performed using 9 independent biological replicates, with 8 plants per replicate. Nitrate content assessment was performed with at least 4 independent biological replicates. Different letters indicate statistical differences between treatments with *p* ≤ 0.05 based on one-way ANOVA Student-Newman-Keuls’ test. The picture represents untreated, treated with K salts, and treated with PSI-362 (50 mg) 12-day-old winter wheat seedlings (cv. Graham) grown in 0 MS agar medium (c).

On the other hand, high levels of K have been documented to improve nitrate uptake (Xu *et al.*, 2020; Fang *et al.*, 2021) and since K content measured in PSI-362 was estimated as moderate (Table 4), additional tests were performed to elucidate any potential effect of this macronutrient on the MOA of PSI-362. In this case, 6-day-old winter wheat seedlings were grown in 0 MS medium supplemented with an equivalent amount of total K present in 50 mg of PSI-362 (Fig. 5c). Plants treated with K_2_SO_4_ and KCl showed a statistically significant lower total plant biomass increase compared to plants treated with PSI-362 in the same medium (−25.6% and −30.6%, respectively) (*p* ≤ 0.001) (Fig. 5a). While plants supplemented with K_2_SO_4_ and KCl showed lower nitrate content than control (−14.1% and −8.2%, respectively), this nitrate depletion was significantly lower than that observed in plants treated with PSI-362 (*p* ≤ 0.001) (Fig. 5b).

### 3.5. PSI-362 affects expression level of genes related to N uptake, transport, and assimilation

PSI-362 has been demonstrated to improve biomass accumulation and nitrate uptake under different nutrient conditions. To gain further insight into its role in enhancing NUE in wheat the differential expression of several key genes, at different time points, PSI-362 dose rate and N supplementation in the growth medium was analysed. The transcript levels of *NITRATE TRANSPORTER1 (TaNRT1.1/NPF6.3),* a dual affinity transceptor involved in nitrate uptake from the soil (Buchner *et al.*, 2015), *NITRATE REDUCTASE1* (*TaNR1*) (Buchner *et al.*, 2015), *NITRITE REDUCTASE1* (*TaNiR1*) (Buchner *et al.*, 2015), *GLUTAMATE DEHYDROGENASE2* (*TaGDH2*) (Buchner *et al.*, 2015), *GLUTAMINE SYNTHETASE1* (*TaGS1*) (Buchner *et al.*, 2015), *NADH-GLUTAMINE OXOGLUTARATE AMINOTRANSFERASE* (*TaNADH-GoGAT*) (Li *et al.*, 2020) and *AMMONIUM TRANSPORTER1* (*TaAMT1*) (Li *et al.*, 2017), which are involved in N assimilation, were evaluated..

Firstly, in order to check whether rapid nitrate uptake and enhanced NUE coincides with the dysregulation of relevant genes, their relative expression was measured after 1 hour, 1 day, and 6 days of treatment with 50 mg of PSI-362 (Fig. 6). Short-term stimulation with PSI-362 for 1 hour showed that *TaNPF6.3, TaGDH2,* and *TaNADH-GoGAT* significantly increased their relative gene expression levels by 1.29, 1.31-, and 1.36-fold compared to control (*p* = 0.016, 0.016, and 0.007, respectively) (Fig. 6a). After applying PSI-362 for 1 day, the gene encoding the transceptor TaNPF6.3, a major sensor of nitrate levels in the soil environment, decreased its relative expression by 1.22-fold (*p* = 0.01) (Fig. 6b), which may be an effect of a negative feedback loop due to the measured nitrate accumulation mentioned earlier (+12.2%) (Fig. 3b). In parallel, *TaNR1* and *TaNiR1* genes, encoding enzymes that metabolize nitrate to nitrite and ammonium respectively, increased their relative gene expression level by 1.03- and 1.13-fold. However, only the *TaNiR1* upregulation was statistically significant (*p* = 0.045). On the other hand, *TaGS1, TaNADH-GoGAT,* and ammonium transporter *TaAMT1* showed moderate decreases in relative gene expression (1.18-, 1.26-, 1.13-fold; *p* = 0.018, 0.005, and 0.035, respectively) (Fig. 6b). Finally, after a long-term stimulation with PSI-362 for 6 days, both transporters *TaNPF6.3* and *TaAMT1*, along with the enzyme *TaGDH2* showed a statistically significant downregulation (1.32-, 1.44-, 1.19-fold; *p* = 0.003, ≤ 0.001, and 0.023, respectively). However, *TaNR1* and *TaNiR1* relative gene expression was clearly upregulated after treating wheat seedlings with PSI-362 for 6 days (1.27- and 1.25-fold; *p* = 0.001 and 0.003, respectively) (Fig. 6c).

**Fig. 6.**
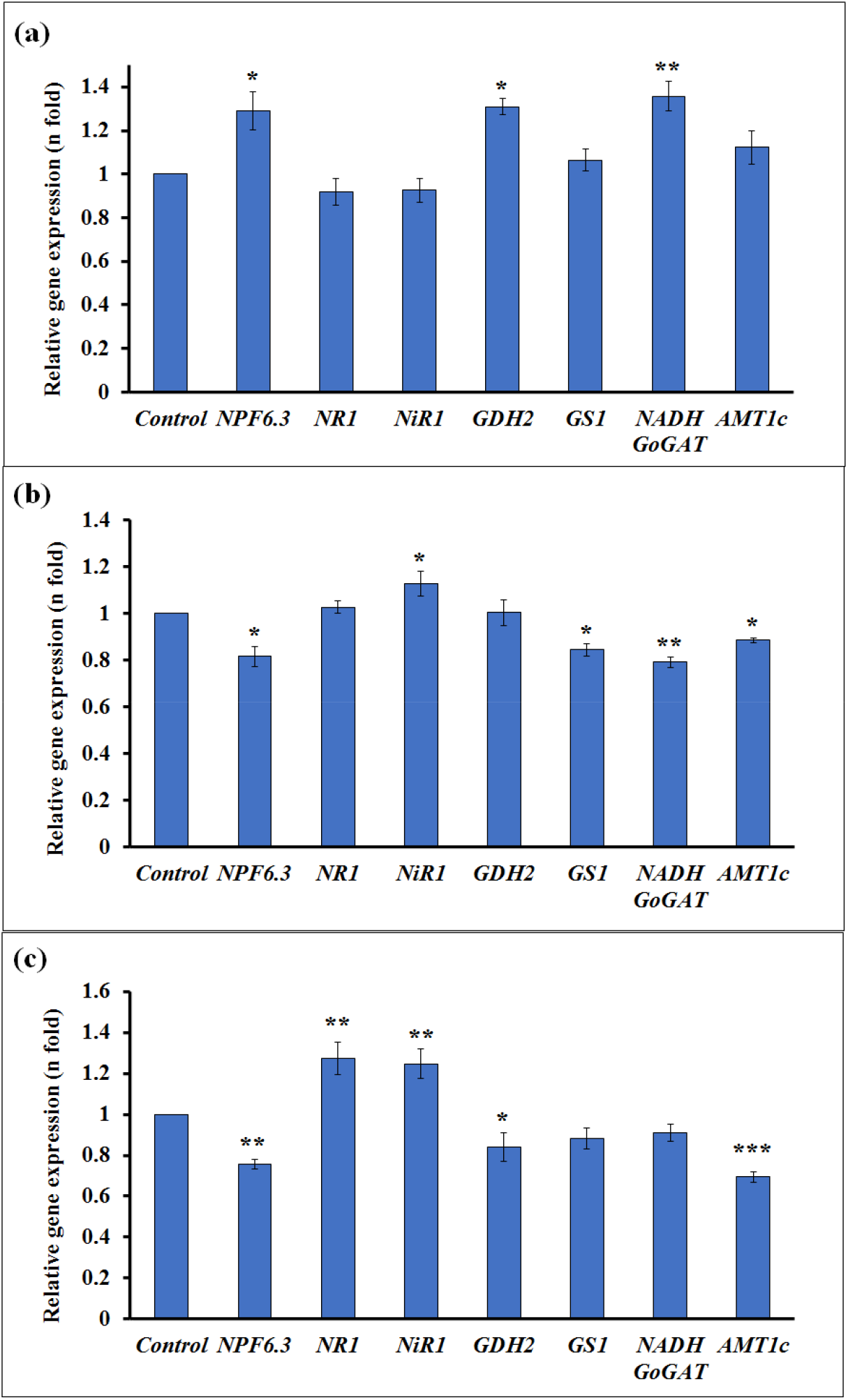
Effects of PSI-362 treatment on relative gene expression in whole wheat seedlings during different time points. 6-day-old winter wheat seedlings were treated with 0 mg (control) and 25 mg PSI-362 in 75ml 1/10 MS agar medium for 1 hour (a), 1 day (b), and 6 days (c). Results are displayed as the relative log2 fold-change with respect to the *TaEF1a* gene expression levels. RNA extractions followed by RT-qPCR experiments were performed with at least 3 independent biological replicates. Means followed by asterisk indicate statistically significant differences between control and PSI-362 treatment within the same application time based on one-way ANOVA Student-Newman-Keuls’ test (***, **, and * significant at *p* ≤ 0.001, *p* ≤ 0.01, and *p* ≤ 0.05, respectively).

The expression of all the genes was also tested after applying a 10-fold lower PSI-362 concentration rate for 6 days. Interestingly, only *TaNPF6.3*, *TaGDH2* and *TaAMT1* were significantly down-regulated between 1.3- and 1.4-fold (*p* = 0.003), and the rest of genes assessed did not show any significant change with respect to control (Fig. S3a). Relative gene expression was also evaluated in plants grown on 10-fold higher N medium (1 MS). A significant upregulation of *TaNR1* (1.31-fold; *p* = 0.02), responsible for nitrate assimilation, and *TaNADH-GoGAT* and *TaAMT1* (1.20- and 1.26-fold; *p* = 0.046 and 0.043, respectively), directly regulating ammonium levels was observed. In parallel, a non-statistically significant down-regulation of *TaGS1* gene, which is involved in glutamine synthesis, was also measured (1.26-fold; *p* = 0.128) (Fig. S3b). These gene expression results highlight how application time, dose and N content in growth medium have a differential effect on PSI-362 impact on multiple genes involved in N uptake, transport, and assimilation.

### 3.6. PSI-362 enhances NUE through N assimilation

Increased biomass, nitrate accumulation and dysregulation of genes related to N uptake, transport and assimilation provides a strong foundation to claim that PSI-362 biostimulant is indeed improving NUE in wheat. However, direct measurements of nitrate assimilation products, amino acids, soluble protein, and photosynthetic pigments content can provide a more detailed insight and understanding of biological events in space and time at metabolic level.

The assessment of N metabolic markers revealed a significant effect of the applied PSI-362 dose rate on wheat plants growing under reduced N (Table 1). Ammonium content showed the opposite trend to that observed with nitrate, decreasing progressively by 10.7% at 5 mg PSI-362 and by 40% at 50 mg dose added to the growth medium (*p* ≤ 0.001). However, the application of PSI-362 did not have any statistically significant effect on total free amino acid (FAA) content at the lowest and highest concentration. Glutamate and glutamine represented nearly 30% of total free amino acids in wheat seedlings grown in our experimental system, therefore measurement of those molecules is relevant to assess the impact of PSI-362 on NUE. Glutamate content increased by 16.1% (*p* = 0.01) and 43.3% (*p* ≤ 0.001) when the lowest and highest PSI-362 dose was applied. However, glutamine content decreased by 12.7 and 26.1% at the same concentrations of PSI-362 (*p* ≤ 0.001). Interestingly, both low and high dose of applied PSI-362 (5 mg and 50 mg) led to elevated accumulation of soluble protein versus control (17.1% and 12.0%; *p* = 0.005 and 0.024, respectively). The accumulation pattern of both chlorophylls (a + b) and carotenoids was similar, increasing by 11.2-15.1% and 13.6-20.0% respectively, after applying 5 and 50 mg of PSI-362 (Table 1).

The short-term treatment (1 hour) with 50 mg PSI-362 under reduced N rate (1/10 MS) did stimulate slight decreases in ammonium and glutamine content (−3.6% and −8.9%; *p* = 0.696 and 0.329, respectively) and a minor glutamate accumulation (+5.1%; *p* = 0.278) (Table S5). PSI-362 application for 1 day resulted in a notable accumulation of several N derived metabolites such as total FAA (+8.9%; *p* = 0.078), glutamate (+33.9%; *p* ≤ 0.001), glutamine (+12.7%; *p* = 0.006), soluble protein (+8.1%; *p* = 0.005) and chlorophylls (a+b) (+14.5%; *p* = 0.012). Long-term treatment with PSI-362 for 6 days led to a significant accumulation of glutamate (+15.7%; *p* = 0.004), soluble protein (13.1%; *p* = 0.005), chlorophylls (a+b) and carotenoids (+15.1-22.1%; *p* ≤ 0.01) along with reduced content of glutamine (−23.6%; *p* = 0.023) and ammonium (−53.2%; *p* ≤ 0.001) compared to control (Table S5).

Regarding the interaction between N rate in growth medium and the biostimulant effect of PSI-362, there were more obvious fluctuations at metabolic than transcriptional level (Table S6, Fig. S3b). Treated wheat seedlings grown for 6 days on 1 MS had significant reductions in glutamate (−17.3%; *p* = 0.018), glutamine (−11,1%; *p* = 0.003) and total FAA content (−11.8%; *p* = 0.099), while ammonium accumulated by 29.6% (*p* ≤ 0.001). However, soluble protein, chlorophylls (a+b) and carotenoids content did not show any significant alteration with respect to control (Table S6). Overall, the results described above indicate that PSI-362 has an effect on each step of N uptake, transport, and assimilation at multiple points, demonstrating an association between phenotype, differential gene expression data and biochemistry.

### 3.7. PSI-362 composition significantly differs from other commercial ANE biostimulants

The commercial ANE biostimulants used in this study showed statistically significant differences in their concentration of macronutrients and the levels of some key components such as ash, seaweed carbohydrates and polyphenols (Table 4). The N:P:K macronutrient content of PSI-362, ANE X and ANE Y was considered very low, with PSI-362 and ANE Y having a total K content above 1% w/v. Overall, the three ANE biostimulants used were primarily composed of ash, uronic acids (representing mainly alginate), fucose, laminarin, free mannitol and polyphenols. However, every formulation had a distinctive signature based on these compositional markers expressed on a dry weight basis.

### 3.8. PSI-362 outperforms other ANE biostimulants enhancing NUE in wheat

The efficacy of applying 50 mg of each commercial ANE for 6 days to enhance NUE in winter wheat seedlings grown on reduced N rate (1/10 MS) was tested using a diverse number of phenotypical and biochemical parameters (Table 5). Total plant, shoot and root biomass in plants treated with PSI-362 (66%, 25%, 93%, respectively) (*p* ≤ 0.001) contrasted with the minor effect obtained for the other biostimulants. Plants treated with the cold extract, ANE X only showed a slight non statistically significant plant biomass increase (+4.2%; *p* = 0.601), characterised by a minor increase in root growth (+14.4%; *p* = 0.262) and no change in shoot biomass. The application of another hot alkaline extract (ANE Y) did increase plant biomass and NUE but this effect was lower than that observed with PSI-362 (+20.6%; *p* = 0.014). The stimulating effect of ANE Y was more noticeable in root biomass (+51.0%; *p* = 0.002) than in shoot tissue (+15.0%; *p* = 0.016) (Table 5). The positive effect of PSI-362 on nitrate uptake (+37.7%; *p* = 0.002) was not observed for treatments ANE X and ANE Y, which only displayed small non-statistically significant increases (+9.4% and +23.4%; *p* = 0.099 and 0.277, respectively) (Table 5). Differences observed in N uptake between applied ANEs were also measured in N assimilation parameters. The consistent accumulation of soluble protein stimulated by PSI-362 (+16.3%) was absent in plants treated with ANE X and ANE Y (−6.9% and +3.3%, respectively). Accumulation of both chlorophylls (a + b) and carotenoids was found in plants treated with PSI-362 (+9.8% and +14.3%, respectively). Remarkably, ANE X had a negligible effect on both photosynthetic pigments content, while ANE Y biostimulant induced a statistically significant decrease of chlorophylls (a + b) and carotenoids by −10.4% and −17.0% (*p* = 0.011 and 0.004, respectively).

**Table 5.**
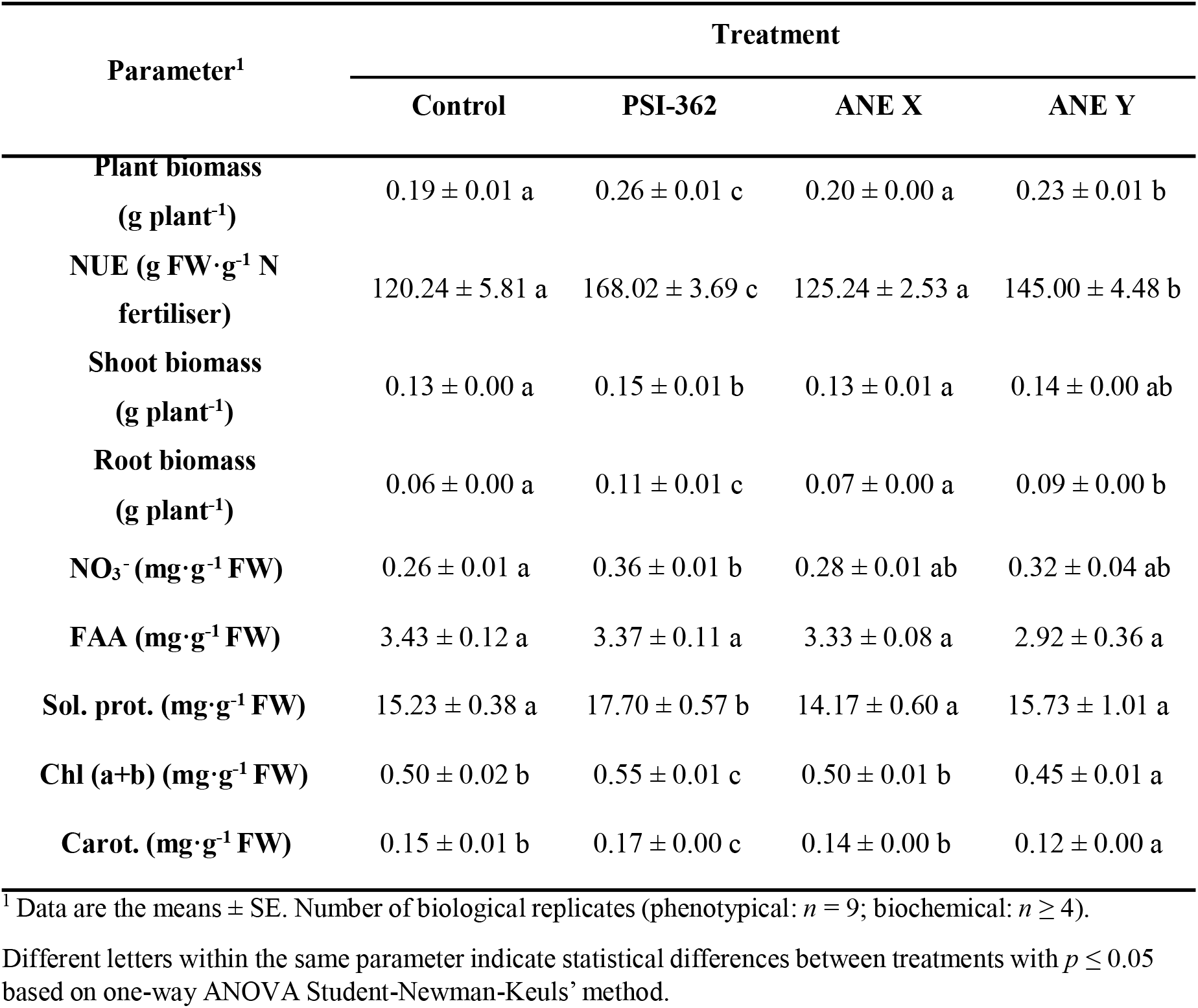
Effect of different ANEs applied for 6 days on phenotypic and biochemical parameters in winter wheat seedlings (cv. Graham) grown on reduced N media (1/10 MS).

## 4. DISCUSSION

NPK are the key macronutrients necessary for plant growth, however, current attention on environmental pollution and climate change has brought their use in agriculture under the spotlight. It has been recently demonstrated that an engineered biostimulant can be used as an enhancer of NUE in barley grown under reduced N supplementation without a yield penalty (Goñi *et al.*, 2021). In the current work we investigated the potential of PSI-362 to improve NUE in another strategic crop, winter wheat. Wheat plants were grown in an enclosed system to gain a better understanding of MOA and growth dynamics (Fig. 1).

### 4.1. PSI-362 promotes biomass increase, however it triggers nitrate uptake before root growth

Plant growth and development is highly dynamic and responsive to environmental stimuli. Availability of macronutrients such as N dictates root growth in order to optimise the uptake in a heterogenous soil. N abundance and starvation typically inhibit root growth, however mild deprivation stimulates root growth to acquire the necessary N (Gruber *et al.*, 2013; Araya *et al.*, 2014). PSI-362 supplementation promoted remarkable biomass production (Fig. 1-3), however in comparison to shoot biomass, root biomass increased more rapidly. Data collected 24h after treatment showing no difference in biomass yet a prominent increase in nitrate content with respect to the control plants, suggests that PSI-362 is most likely increasing the capacity of existing channels before producing more root surface (Fig. 3).

PSI-362 promotes growth and specifically root growth not only in N deficient media, but also in media saturated with N (Fig. 2). It indicates that PSI-362 stimulation overwrites the canonical plant root growth response to N (O’Brien *et al.*, 2016), breaking the negative feedback loop related to high N metabolite accumulation. Moreover, wheat seedlings stimulated with PSI-362 and grown on deficient and replete media reached a similar biomass accumulation, indicating the limit of enhancing NUE.

### 4.2. PSI-362 induces the accumulation of nitrate over a period of time

N availability is the main environmental cue for plant growth and development. Despite the comprehensive regulation of nitrate sensing, transport and signalling at local level, nitrate is a long-distance signalling molecule, conveying the N status at whole-plant level (Araya *et al.*, 2014; Ota *et al.*, 2020; Fang *et al.*, 2021). Therefore, measuring nitrate content is one of the key parameters to assess the efficacy of NUE in plants. PSI-362 treatments independently of the applied concentration and N status in the MS medium promoted significant increase of nitrate content (Fig. 1-3). The measurements of nitrate content in the 1/10 MS medium at different time points after treatment (1, 3, 6, and 9 days) clearly demonstrated the ability of PSI-362 to induce rapid uptake until the resources get depleted (Fig. 4). The experimental setup utilised provides evidence that PSI-362 induces a rapid uptake of N, at different N levels and dose rates without a phytotoxic effect and does this over a prolonged period of time.

### 4.3. PSI-362 dysregulates major genetic markers of N uptake and assimilation

Plants, as sessile organisms adjust their growth and development according to external cues. Specific growth modifications are determined by the type of the cue and response time, usually first in the form of transcription and subsequently translation followed by physiological and biochemical alterations. In order to understand how PSI-362 triggers NUE in wheat, several crucial genetic markers were tested. While *NRT1.1/NPF6.3* is canonically up-regulated in an environment deficient in N (Loqué *et al.*, 2003), PSI-362 short-term application for 1 hour induced the expression even further than the control plants along with two other genes that encode enzymes that assimilate ammonium in the form of glutamate (*TaGDH2* and *TaGoGAT*) (Fig. 6). It indicates that the treated wheat plant is creating resources for higher N uptake and assimilation. However, *TaNPF6.3* was already down-regulated 24 hours after treatment, which is likely associated with a negative feedback loop because of higher nitrate content in the plant tissue. Such correlations between gene expression and nitrate content were observed in all long-term treated plants (6 days), independently of PSI-362 concentration or N supplementation (Fig. 6, Fig. S3). Low expression of the transceptor in the context of high nitrate accumulation is not surprising, however it is intriguing, when taking into account the remarkable root growth in treated plants (Fig. 1-3) (Loqué *et al.*, 2003). Prolonged PSI-362 treatment (6 days) altered the plant developmental program by changing the nutrients status in the enclosed system, therefore the interpretation of gene expression data can be challenging due to multiple genetic interactions that have already occurred. Unfortunately, the perception and signalling mechanism of PSI-362 complex remains unknown. However, taking into consideration the remarkable root growth in high N medium, nitrate accumulation, glutamate content increase, and upregulation of *TaGDH2* and *TaGoGAT* genes in the first hour of the stimulation, it seems that the family of glutamate receptors-like GLRs (iGluR-related) could be a possible candidate to antagonise *TaNRT1.1/NPF6.3* action and promote root growth. Canonically, root growth is inhibited in the presence of abundant nitrate or amino acids, however glutamate is known to elicit changes in growth and root branching (Forde, 2014). It is documented that GLRs, that operate as non-specific cation channels (NSCC), can release Ca^2+^ from intracellular stores to alter gene transcription in response to intracellular amino acid levels (Davenport, 2002; Vincill *et al.*, 2013). In this context, further investigation of GLRs family or *MEKK1* genes, a positive regulator of glutamate signalling would be particularly interesting (Davenport, 2002). Furthermore, a direct link between N status and cell cycle progression is attributed to TEOSINT BRANCHED1/CYCLOIDEA/PROLIFERATING CELL FACTOR1-20 (TCP20) – NIN-LIKE PROTEIN 6/7 (NLP6/7) regulatory nexus (Guan, 2017). TCP functions as the main regulators of plant morphology with an emerging role as integrators of plant developmental responses to the environment. This is especially true in root meristem growth that is regulated by intertwined nitrate, phytohormones and glucose-TOR signalling pathway (Guan, 2017; McCready *et al.*, 2020). In order to obtain full spatiotemporal information on phenotypic, biochemical, and genetic interplay, a full transcript sequencing at several time points would be critical.

### 4.4. PSI-362 affects every step of N assimilation including protein synthesis

In order to gain a better understanding of the nature of PSI-362 stimulation at phenotypic and genetic level, we measured a number of biochemical parameters. PSI-362 application showed a very clear effect on transcript levels after 1 hour of treatment, however biochemical analysis did not show any notable changes except a slight increase in glutamate content, that might be associated with *TaGDH2* and *TaGoGAT* up-regulation (Fig. 6). The early expression peak of both genes might also explain the significant further accumulation of glutamate and glutamine along with reduced ammonium content 24 hours after PSI-362 treatment (Table S4). The observed up-regulation of *TaNiR1*, indicated that ammonium produced from nitrate is metabolised very rapid after treatment with PSI-362 (Fig. 6). Six days of PSI-362 treatment decreased glutamine content compared to control. However, the same treatment increased glutamate content and induced low ammonium levels. These results seem contradictory with the up-regulation of *TaNR1* and *TaNiR1* (Fig. 6), which in theory are associated with higher ammonium production rate (Hirel & Krapp, 2020). However, this behaviour also exposes the long-term signalling activity on the activation of ammonium recycling and glutamate biosynthesis through modulation of GS/GOGAT and GDH enzymes (Pratelli & Pilot, 2014). As previously observed in barley grown under real field conditions (Goñi *et al.*, 2021), PSI-362 increased both soluble protein and photosynthetic pigments by 10-15%, with this accumulation being associated with a higher content of direct precursors of these macromolecules such as glutamate (Forde & Lea, 2007). Increased soluble protein content and chlorophylls (a+b) under reduced N conditions suggests that PSI-362 not only impacted downstream steps of N assimilation but also influences carbon fixation and photosynthetic efficiency (Feller *et al.*, 2008; Cabrera-Bosquet *et al.*, 2009).

The biomass, nitrate uptake and gene expression analysis demonstrate that the PSI-362 effect on wheat NUE mechanisms are differing depending on N availability. Biochemical analysis of treated wheat seedlings grown in medium saturated with N (1 MS) showed that they accumulated ammonium with respect to control (Table S5). While high ammonium concentrations are known to be phytotoxic (Hachiya *et al.*, 2021), absolute values were still lower than those observed in wheat grains during grain filling (Wei *et al.*, 2021), which may explain why plants stimulated with PSI-362 under high N rate develop a healthy phenotype (Fig. 2). The observed dysregulation of *TaAMT1* ammonium transporter also correlated with ammonium content change in treated plants grown under reduced and full N rate (Fig. 6, Fig. S3). Interestingly, it has been observed how gene expression levels of *AMT* transporters are sensitive to ammonium content in wheat, rice, and Arabidopsis (Li *et al.*, 2017). Unlike treated plants grown on 1/10 MS, the application of PSI-362 under high N rate reduced plant glutamate content and did not induce any significant change in soluble proteins and photosynthetic pigments (Table S5). This data suggests that the combination of PSI-362 with high N supplementation was less effective in synchronizing N uptake and assimilation biochemical processes, leading to similar biomass gains in treated plants grown on both N rates. This bottleneck in biomass accumulation could be attributed to the inherent genetic limitation of wheat seedling growing in our current system. It is tempting to speculate that this limit could be increased further if photosynthesis and carbon central metabolic pathways, that are source of energy (ATP), reduction intermediates (NADH/NADHP), and indispensable precursors (2-oxoglutarate) for N assimilation, were more efficient (Krapp, 2015).

### 4.5. Comparative analysis of commercial ANEs in context of enhanced NUE

Published literature on the application of ANEs to improve NUE have been mainly focused on acidic extracts or combinations of ANE with amino acids on high N demanding crops under optimum nutrient conditions (Jannin *et al.*, 2013; Billard *et al.*, 2014; Stamatiadis *et al.*, 2015; Laurent *et al.*, 2020). However, there is no relevant information about the comparative effect of commercial ANEs manufactured using different extraction processes in improving NUE in crops under reduced N rate. Overall, we found significant differences in the efficacy of three commercial ANE biostimulants in improving NUE in winter wheat. While PSI-362 consistently increases plant biomass, nitrate content, soluble protein, and photosynthetic pigments content under reduced N rate (Table 5), another hot alkaline extract (ANE Y) was not able to coordinate a significant accumulation of both nitrate and N assimilation products, which translated into significantly reduced gains in NUE. Furthermore, the application of ANE X, manufactured using a proprietary process at low temperatures, did not have any relevant effect in stimulating any NUE related phenotypical or biochemical parameter (Table 5). Although PSI-362 and ANE Y did not contain any significant N content to explain their biostimulant activity, the K content in both extracts may suggest the presence of a positive interaction with N in the 1/10 MS growth medium (Raddatz *et al.*, 2020; Xu *et al.*, 2020). However, K supplementation by its own had only a minor effect in enhancing biomass growth and nitrate uptake of winter wheat seedlings (Fig. 5). While produced from the same raw material, the tested ANE biostimulants were compositionally diverse in terms of ash, carbohydrates, and polyphenol content. This compositional variation has been observed previously and has been associated with biostimulant efficacy in various agronomic productivity challenges (Goni *et al.*, 2016; Ertani *et al.*, 2018; Goñi *et al.*, 2018; Carmody *et al.*, 2020). The results emphasise the role of processing conditions in the production of biostimulants with optimised effectiveness to solve specific agronomic problems.

### 4.6. Summary and perspectives

The demand for technologies that can address the urgent need to reduce global warming is growing rapidly as government policies are being implemented in order to meet international climate change targets. Agriculture is one of the sectors in urgent need to such technologies in order to be able to continue to meet global demand for food while reducing emissions. N is an essential input in all modern, commercially viable, agricultural production systems, yet its loss to the environment needs to be reduced. Plant biostimulants are an emerging regulated technology which have the potential to have a major role in delivering more sustainable agricultural practices. Biostimulants can achieve this by improving productivity when crops are challenged by nutrient deficiencies and abiotic stress. However, their ability to achieve this in practice needs to be supported by strong scientific evidence demonstrating performance that leads to increase productivity or enhanced resource use efficiency. The data presented in this research highlights the need for such scientific rigor, as some products classed as ANE biostimulants do not have any effect on increasing nitrogen use efficiency, while the precision engineered PSI-362 displays potential to have a significant impact in enhancing NUE. The expansion of the applicability in other species of staple crop types such as maize or potatoes depend on the level of understanding of yield production mechanism in relation to N status. The current data does not allow the mechanism to be defined unequivocally, which presumably changes, adapting to the developmental stage of the plant, N status inside the plant and in the environment. To answer that question further testing of phosphorylation status of the transporters and their turnover in the plasma membrane would be interesting to investigate. PSI-362 technology combined with good practices of fertiliser application and field management, may in the near future be a feasible solution to reduce the N consumption.

## 5. AUTHOR CONTRIBUTIONS

Ł.Ł., O.G., and S.O’C. conceived and designed the experiments. Ł.Ł., O.G., E.I., E.F., performed the experiments and analysed the data. Ł.Ł., O.G., and S.O’C. wrote the article. All the authors reviewed and approved the article.

## 6. ACKNOWLEDGEMENTS

The authors would like to thank Brandon Bioscience for the gift of the ANE-based biostimulants (PSI-362, ANE X, and ANE Y) used in this study.

## 9. SUPPLEMENTARY MATERIAL

The following Supporting Material is available for this article:

Fig. S1. PSI-362 enhances plant growth in winter wheat seedlings during different time points.

Fig. S2. PSI-362 increases plant biomass in winter wheat seedlings grown in nutrient starving conditions

Fig. S3. Changes in relative gene expression in whole wheat seedlings with low PSI-362 concentration rate or using high N growth medium

Table S1. Primers used for RT-qPCR analysis in winter wheat (cv. Graham)

Table S2. Effect of PSI-362 concentration applied for 6 days on NUE in winter wheat seedlings (cv. Graham) grown on reduced N media (1/10 MS).

Table S3 Effect of PSI-362 applied for 6 days on NUE in winter wheat seedlings (cv. Graham) grown on reduced (1/10 MS) and saturated N media (1 MS).

Table S4. Effect of PSI-362 applied during different time points on NUE in winter wheat seedlings (cv. Graham) grown on reduced N media (1/10 MS).

Table S5. Analysis of variance and mean comparisons for biochemical parameters in winter wheat seedlings (cv. Graham) grown on reduced N media (1/10 MS) and treated with PSI-362 during different time points.

Table S6. Analysis of variance and mean comparisons for biochemical parameters in winter wheat seedlings (cv. Graham) treated with PSI-362 for 6 days and growing under two different N rates.

## Abbreviations

ANE: *Ascophyllum nodosum* extract
DEEMM: diethylethoxymethylenemalonate
FW: fresh weight
Fd: ferrodoxin
FAA: free amino acids
GHG: greenhouse gas
Glu: glutamate
Gln: glutamine
ICP-MS: inductively coupled plasma mass spectrometry
MOA: mode of action
NO_3_-: MS medium, Murashige and Skoog medium
NO_3_-: nitrate
NH_4_+: ammonium
NUE: nitrogen use efficiency
RH: relative humidity
RP-HPLC: reverse phase high performance liquid chromatography
TCA: tricarboxylic acid cycle

## REFERENCES

Araya T, Miyamoto M, Wibowo J, Suzuki A, Kojima S, Tsuchiya YN, Sawa S, Fukuda H, von Wirén N, Takahashi H. 2014. CLE-CLAVATA1 peptide-receptor signaling module regulates the expansion of plant root systems in a nitrogen-dependent manner. Proceedings of the National Academy of Sciences 111(5): 2029.

Bajgain P, Russell B, Mohammadi M. 2018. Phylogenetic analyses and in-seedling expression of ammonium and nitrate transporters in wheat. Scientific Reports 8(1): 7082.

Beig B, Niazi MBK, Jahan Z, Hussain A, Zia MH, Mehran MT. 2020. Coating materials for slow release of nitrogen from urea fertilizer: a review. Journal of Plant Nutrition 43(10): 1510–1533.

Bellegarde F, Gojon A, Martin A. 2017. Signals and players in the transcriptional regulation of root responses by local and systemic N signaling in Arabidopsis thaliana. J Exp Bot 68(10): 2553–2565.

Billard V, Etienne P, Jannin L, Garnica M, Cruz F, Garcia-Mina J-M, Yvin J-C, Ourry A. 2014. Two Biostimulants Derived from Algae or Humic Acid Induce Similar Responses in the Mineral Content and Gene Expression of Winter Oilseed Rape (Brassica napus L.). Journal of Plant Growth Regulation 33(2): 305–316.

Bouguyon E, Perrine-Walker F, Pervent M, Rochette J, Cuesta C, Benkova E, Martinière A, Bach L, Krouk G, Gojon A, et al. 2016. Nitrate Controls Root Development through Posttranscriptional Regulation of the NRT1.1/NPF6.3 Transporter/Sensor. Plant physiology 172(2): 1237–1248.

Buchner P, Tausz M, Ford R, Leo A, Fitzgerald GJ, Hawkesford MJ, Tausz-Posch S. 2015. Expression patterns of C- and N-metabolism related genes in wheat are changed during senescence under elevated CO2 in dry-land agriculture. Plant Sci 236: 239–249.

Cabrera-Bosquet L, Albrizio R, Araus JL, Nogués S. 2009. Photosynthetic capacity of field-grown durum wheat under different N availabilities: A comparative study from leaf to canopy. Environmental and Experimental Botany 67(1): 145–152.

Carmody N, Goñi O, Łangowski Ł, O’Connell S. 2020. Ascophyllum nodosum Extract Biostimulant Processing and Its Impact on Enhancing Heat Stress Tolerance During Tomato Fruit Set. Frontiers in plant science 11: 807–807.

Chamizo-Ampudia A, Sanz-Luque E, Llamas A, Galvan A, Fernandez E. 2017. Nitrate Reductase Regulates Plant Nitric Oxide Homeostasis. Trends in Plant Science 22(2): 163–174.

Chislock MF, Sharp KL, Wilson AE. 2014. Cylindrospermopsis raciborskii dominates under very low and high nitrogen-to-phosphorus ratios. Water Research 49: 207–214.

Davenport R. 2002. Glutamate Receptors in Plants. Annals of Botany 90(5): 549–557.

Dimkpa CO, Fugice J, Singh U, Lewis TD. 2020. Development of fertilizers for enhanced nitrogen use efficiency – Trends and perspectives. Science of The Total Environment 731: 139113.

du Jardin P. 2015. Plant biostimulants: Definition, concept, main categories and regulation. Scientia Horticulturae 196: 3–14.

Efretuei A, Gooding M, White EM, Spink J, Hackett R. 2016. Effect of nitrogen fertilizer application timing on nitrogen use efficiency and grain yield of winter wheat in Ireland.

Ertani A, Francioso O, Tinti A, Schiavon M, Pizzeghello D, Nardi S. 2018. Evaluation of Seaweed Extracts From Laminaria and Ascophyllum nodosum spp. as Biostimulants in Zea mays L. Using a Combination of Chemical, Biochemical and Morphological Approaches. Front Plant Sci 9: 428.

Fang XZ, Fang SQ, Ye ZQ, Liu D, Zhao KL, Jin CW. 2021. NRT1.1 Dual-Affinity Nitrate Transport/Signalling and its Roles in Plant Abiotic Stress Resistance. Frontiers in plant science 12(1817).

Feller U, Anders I, Mae T. 2008. Rubiscolytics: fate of Rubisco after its enzymatic function in a cell is terminated. J Exp Bot 59(7): 1615–1624.

Forde BG. 2014. Nitrogen signalling pathways shaping root system architecture: an update. Current Opinion in Plant Biology 21: 30–36.

Forde BG, Lea PJ. 2007. Glutamate in plants: metabolism, regulation, and signalling. Journal of Experimental Botany 58(9): 2339–2358.

Foyer CH, Noctor G. 2011. Ascorbate and glutathione: the heart of the redox hub. Plant Physiol 155(1): 2–18.

Frioni T, Sabbatini P, Tombesi S, Norrie J, Poni S, Gatti M, Palliotti A. 2018. Effects of a biostimulant derived from the brown seaweed Ascophyllum nodosum on ripening dynamics and fruit quality of grapevines. Scientia Horticulturae 232: 97–106.

Fu Y-F, Zhang Z-W, Yang X-Y, Wang C-Q, Lan T, Tang X-Y, Chen G-D, Zeng J, Yuan S. 2020. Nitrate reductase is a key enzyme responsible for nitrogen-regulated auxin accumulation in Arabidopsis roots. Biochemical and Biophysical Research Communications 532(4): 633–639.

Gaudinier A, Rodriguez-Medina J, Zhang L, Olson A, Liseron-Monfils C, Bågman, A-M, Foret J, Abbitt S, Tang M, Li B, et al. 2018. Transcriptional regulation of nitrogen-associated metabolism and growth. Nature 563(7730): 259–264.

Gil-Ortiz R, Naranjo MÁ, Ruiz-Navarro A, Caballero-Molada M, Atares S, García C, Vicente O. 2020. New Eco-Friendly Polymeric-Coated Urea Fertilizers Enhanced Crop Yield in Wheat. Agronomy 10(3): 438.

Gómez-Alonso S, Hermosín-Gutiérrez I, García-Romero E. 2007. Simultaneous HPLC analysis of biogenic amines, amino acids, and ammonium ion as aminoenone derivatives in wine and beer samples. J Agric Food Chem 55(3): 608–613.

Goni O, Fort A, Quille P, McKeown PC, Spillane C, O’Connell S. 2016. Comparative Transcriptome Analysis of Two Ascophyllum nodosum Extract Biostimulants: Same Seaweed but Different. J Agric Food Chem 64(14): 2980–2989.

Goñi O, Łangowski Ł, Feeney E, Quille P, O’Connell S. 2021. Reducing Nitrogen Input in Barley Crops While Maintaining Yields Using an Engineered Biostimulant Derived From Ascophyllum nodosum to Enhance Nitrogen Use Efficiency. Front Plant Sci 12: 664682.

Goñi O, Quille P, O’Connell S. 2018. Ascophyllum nodosum extract biostimulants and their role in enhancing tolerance to drought stress in tomato plants. Plant Physiol Biochem 126: 63–73.

Gruber BD, Giehl RFH, Friedel S, von Wirén N. 2013. Plasticity of the Arabidopsis Root System under Nutrient Deficiencies Plant physiology 163(1): 161–179.

Grzechowiak M, Sliwiak J, Jaskolski M, Ruszkowski M. 2020. Structural Studies of Glutamate Dehydrogenase (Isoform 1) From Arabidopsis thaliana, an Important Enzyme at the Branch-Point Between Carbon and Nitrogen Metabolism. Frontiers in plant science 11(754).

Guan P. 2017. Dancing with Hormones: A Current Perspective of Nitrate Signaling and Regulation in Arabidopsis. Frontiers in plant science 8(1697).

Gutiérrez Rodrigo A. 2012. Systems Biology for Enhanced Plant Nitrogen Nutrition. Science 336(6089): 1673–1675.

Hachiya T, Inaba J, Wakazaki M, Sato M, Toyooka K, Miyagi A, Kawai-Yamada M, Sugiura D, Nakagawa T, Kiba T, et al. 2021. Excessive ammonium assimilation by plastidic glutamine synthetase causes ammonium toxicity in Arabidopsis thaliana. Nature Communications 12(1): 4944.

He X, Qu B, Li W, Zhao X, Teng W, Ma W, Ren Y, Li B, Li Z, Tong Y. 2015. The Nitrate-Inducible NAC Transcription Factor TaNAC2-5A Controls Nitrate Response and Increases Wheat Yield Plant physiology 169(3): 1991–2005.

Hirel B, Krapp A 2020. Nitrogen utilization in plants I biological and agronomic importance. Encyclopedia of Biochemistry. 3rd Edition. Elsevier.

Jannin L, Arkoun M, Etienne P, Laîné P, Goux D, Garnica M, Fuentes M, Francisco SS, Baigorri R, Cruz F, et al. 2013. Brassica napus Growth is Promoted by Ascophyllum nodosum (L.) Le Jol.Seaweed Extract: Microarray Analysis and Physiological Characterization of N, C, and S Metabolisms. Journal of Plant Growth Regulation 32(1): 31–52.

Kanter DR, Zhang X, Mauzerall DL. 2015. Reducing nitrogen pollution while decreasing farmers’ costs and increasing fertilizer industry profits. Journal of environmental quality 44(2): 325–335.

Kiba T, Krapp A. 2016. Plant Nitrogen Acquisition Under Low Availability: Regulation of Uptake and Root Architecture. Plant Cell Physiol 57(4): 707–714.

Konishi N, Ishiyama K, Matsuoka K, Maru I, Hayakawa T, Yamaya T, Kojima S. 2014. NADH-dependent glutamate synthase plays a crucial role in assimilating ammonium in the Arabidopsis root. Physiol Plant 152(1): 138–151.

Krapp A. 2015. Plant nitrogen assimilation and its regulation: a complex puzzle with missing pieces. Curr Opin Plant Biol 25: 115–122.

Łangowski Ł, Goñi O, Marques FS, Hamawaki OT, da Silva CO, Nogueira APO, Teixeira MAJ, Glasenapp JS, Pereira M, O’Connell S. 2021. Ascophyllum nodosum Extract (SealicitTM) Boosts Soybean Yield Through Reduction of Pod Shattering-Related Seed Loss and Enhanced Seed Production. Frontiers in plant science 12(176).

Łangowski Ł, Goñi O, Quille P, Stephenson P, Carmody N, Feeney E, Barton D, Østergaard L, O’Connell S. 2019. A plant biostimulant from the seaweed Ascophyllum nodosum (Sealicit) reduces podshatter and yield loss in oilseed rape through modulation of IND expression. Scientific Reports 9(1): 16644.

Lassaletta L, Billen, G., Garnier, J., Bouwman, L., Velazquez, E., Mueller, N. D., & Gerber, J. S. 2016. Nitrogen use in the global food system: Past trends and future trajectories of agronomic performance, pollution, trade, and dietary demand. Environmental Research Letters 11(9).

Laurent E-A, Ahmed N, Durieu C, Grieu P, Lamaze T. 2020. Marine and fungal biostimulants improve grain yield, nitrogen absorption and allocation in durum wheat plants. The Journal of Agricultural Science 158(4): 279–287.

Li S, Zhang C, Li J, Yan L, Wang N, Xia L. 2021. Present and future prospects for wheat improvement through genome editing and advanced technologies. Plant Communications 2(4): 100211.

Li T, Liao K, Xu X, Gao Y, Wang Z, Zhu X, Jia B, Xuan Y. 2017. Wheat Ammonium Transporter (AMT) Gene Family: Diversity and Possible Role in Host–Pathogen Interaction with Stem Rust. Frontiers in plant science 8(1637).

Li W, He X, Chen Y, Jing Y, Shen C, Yang J, Teng W, Zhao X, Hu W, Hu M, et al. 2020. A wheat transcription factor positively sets seed vigour by regulating the grain nitrate signal. New Phytologist 225(4): 1667–1680.

Liao M, Fillery IRP, Palta JA. 2004. Early vigorous growth is a major factor influencing nitrogen uptake in wheat. Funct Plant Biol 31(2): 121–129.

Liu Y, Wang C, He N, Wen X, Gao Y, Li S, Niu S, Butterbach-Bahl K, Luo Y, Yu G. 2017. A global synthesis of the rate and temperature sensitivity of soil nitrogen mineralization: latitudinal patterns and mechanisms. Global Change Biology 23(1): 455–464.

Loqué D, Tillard P, Gojon A, Lepetit M. 2003. Gene expression of the NO3-transporter NRT1.1 and the nitrate reductase NIA1 is repressed in Arabidopsis roots by NO2-, the product of NO3-reduction. Plant physiology 132(2): 958–967.

Maia LB, Moura JJG. 2014. How Biology Handles Nitrite. Chemical Reviews 114(10): 5273–5357.

Masclaux-Daubresse C, Daniel-Vedele F, Dechorgnat J, Chardon F, Gaufichon L, Suzuki A. 2010. Nitrogen uptake, assimilation and remobilization in plants: challenges for sustainable and productive agriculture. Ann Bot 105(7): 1141–1157.

McCready K, Spencer V, Kim M. 2020. The Importance of TOR Kinase in Plant Development. Front Plant Sci 11: 16.

Miller A, Cramer M. 2005. Root nitrogen acquisition and assimilation. Plant and Soil 274(1): 1–36.

Miller AJ, Cramer MD. 2005. Root Nitrogen Acquisition and Assimilation. Plant and Soil 274(1): 1–36.

O’Brien José A, Vega A, Bouguyon E, Krouk G, Gojon A, Coruzzi G, Gutiérrez Rodrigo A. 2016. Nitrate Transport, Sensing, and Responses in Plants. Molecular Plant 9(6): 837–856.

Ota R, Ohkubo Y, Yamashita Y, Ogawa-Ohnishi M, Matsubayashi Y. 2020. Shoot-to-root mobile CEPD-like 2 integrates shoot nitrogen status to systemically regulate nitrate uptake in Arabidopsis. Nature Communications 11(1): 641.

Pfromm PH. 2017. Towards sustainable agriculture: Fossil-free ammonia. Journal of Renewable and Sustainable Energy 9(3): 034702.

Pratelli R, Pilot G. 2014. Regulation of amino acid metabolic enzymes and transporters in plants. J Exp Bot 65(19): 5535–5556.

Raddatz N, Morales de Los Ríos L, Lindahl M, Quintero FJ, Pardo JM. 2020. Coordinated Transport of Nitrate, Potassium, and Sodium. Frontiers in plant science 11: 247–247.

Raun WR, Johnson GV. 1999. Improving Nitrogen Use Efficiency for Cereal Production. Agronomy Journal 91(3): 357–363.

Ristova D, Carré C, Pervent M, Medici A, Kim GJ, Scalia D, Ruffel S, Birnbaum KD, Lacombe B, Busch W, et al. 2016. Combinatorial interaction network of transcriptomic and phenotypic responses to nitrogen and hormones in the Arabidopsis thaliana root. Sci Signal 9(451): rs13.

Rouphael Y, Colla G. 2020. Editorial: Biostimulants in Agriculture. Frontiers in plant science 11(40).

Sandhu N, Sethi M, Kumar A, Dang D, Singh J, Chhuneja P. 2021. Biochemical and Genetic Approaches Improving Nitrogen Use Efficiency in Cereal Crops: A Review. Frontiers in plant science 12(757).

Schaufler G, Kitzler B, Schindlbacher A, Skiba U, Sutton MA, Zechmeister-Boltenstern S. 2010. Greenhouse gas emissions from European soils under different land use: effects of soil moisture and temperature. European Journal of Soil Science 61(5): 683–696.

Shukla PS, Mantin EG, Adil M, Bajpai S, Critchley AT, Prithiviraj B. 2019. Ascophyllum nodosum-Based Biostimulants: Sustainable Applications in Agriculture for the Stimulation of Plant Growth, Stress Tolerance, and Disease Management. Frontiers in plant science 10(655).

Snyder CS. 2017. Enhanced nitrogen fertiliser technologies support the ‘4R’ concept to optimise crop production and minimise environmental losses. Soil Research 55(6): 463–472.

Stamatiadis S, Evangelou L, Yvin J-C, Tsadilas C, Mina JMG, Cruz F. 2015. Responses of winter wheat to Ascophyllum nodosum (L.) Le Jol. extract application under the effect of N fertilization and water supply. Journal of Applied Phycology 27(1): 589–600.

Tercé-Laforgue T, Bedu M, Dargel-Grafin C, Dubois F, Gibon Y, Restivo FM, Hirel B. 2013. Resolving the Role of Plant Glutamate Dehydrogenase: II. Physiological Characterization of Plants Overexpressing the Two Enzyme Subunits Individually or Simultaneously. Plant and Cell Physiology 54(10): 1635–1647.

TransparencyMarketResearch. 2021. Biostimulants Market - Global Industry Analysis, Size, Share, Growth, Trends, and Forecast, 2021-2031: Transparency Market Research.

Vance CP. 2001. Symbiotic nitrogen fixation and phosphorus acquisition. Plant nutrition in a world of declining renewable resources. Plant Physiol 127(2): 390–397.

Vidal EA, Alvarez JM, Araus V, Riveras E, Brooks MD, Krouk G, Ruffel S, Lejay L, Crawford NM, Coruzzi GM, et al. 2020. Nitrate in 2020: Thirty Years from Transport to Signaling Networks. Plant Cell 32(7): 2094–2119.

Vincill ED, Clarin AE, Molenda JN, Spalding EP. 2013. Interacting Glutamate Receptor-Like Proteins in Phloem Regulate Lateral Root Initiation in Arabidopsis The Plant Cell 25(4): 1304–1313.

Wang F, Liu Yp, Zhang H, Chu K. 2019. CuO/graphene nanocomposite for nitrogen reduction reaction. ChemCatChem 11(5): 1441–1447.

Wang Q, Nian J, Xie X, Yu H, Zhang J, Bai J, Dong G, Hu J, Bai B, Chen L, et al. 2018. Genetic variations in ARE1 mediate grain yield by modulating nitrogen utilization in rice. Nature Communications 9(1): 735.

Wang Y-Y, Cheng Y-H, Chen K-E, Tsay Y-F. 2018. Nitrate Transport, Signaling, and Use Efficiency. Annual Review of Plant Biology 69(1): 85–122.

Wei Y, Xiong S, Zhang Z, Meng X, Wang L, Zhang X, Yu M, Yu H, Wang X, Ma X. 2021. Localization, Gene Expression, and Functions of Glutamine Synthetase Isozymes in Wheat Grain (Triticum aestivum L.). Frontiers in plant science 12(37).

White PJ, Brown PH. 2010. Plant nutrition for sustainable development and global health. Annals of Botany 105(7): 1073–1080.

Xu X, Du X, Wang F, Sha J, Chen Q, Tian G, Zhu Z, Ge S, Jiang Y. 2020. Effects of Potassium Levels on Plant Growth, Accumulation and Distribution of Carbon, and Nitrate Metabolism in Apple Dwarf Rootstock Seedlings. Front Plant Sci 11: 904.

Yan M, Pan G, Lavallee JM, Conant RT. 2020. Rethinking sources of nitrogen to cereal crops. Global Change Biology 26(1): 191–199.

Yu C, Liu Y, Zhang A, Su S, Yan A, Huang L, Ali I, Liu Y, Forde BG, Gan Y. 2015. MADS-box Transcription Factor OsMADS25 Regulates Root Development through Affection of Nitrate Accumulation in Rice. PLoS One 10(8): e0135196.

Yu T, Zhuang Q. 2020. Modeling biological nitrogen fixation in global natural terrestrial ecosystems. Biogeosciences 17(13): 3643–3657.

Zörb C, Ludewig U, Hawkesford MJ. 2018. Perspective on Wheat Yield and Quality with Reduced Nitrogen Supply. Trends in Plant Science 23(11): 1029–1037.

