## Supplementary data for "A mechanistic investigation of enhanced nitrogen use efficiency in wheat seedlings after treatment with an *Ascophyllum nodosum* biostimulant"

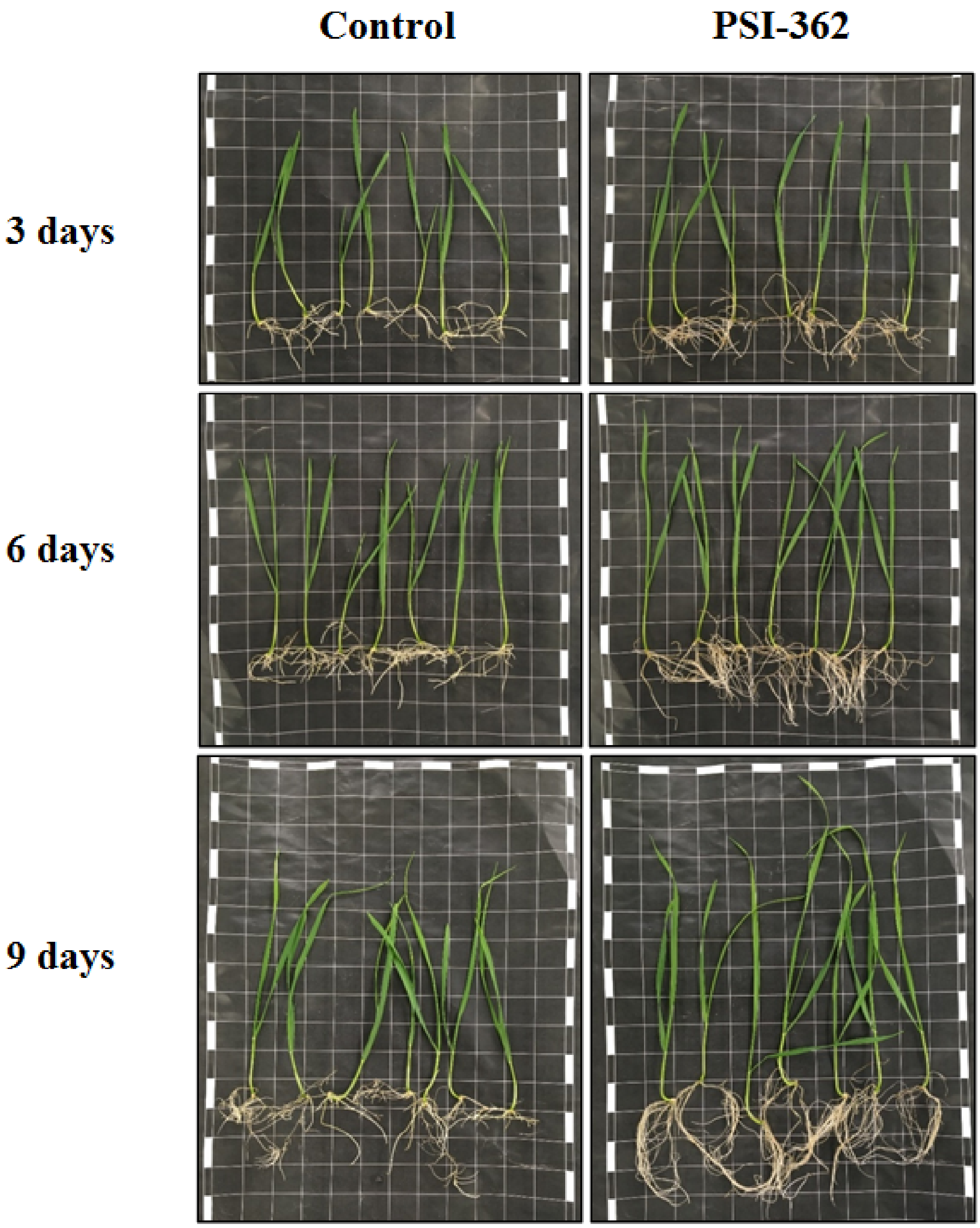


**Fig. S1.** PSI-362 enhances plant growth in winter wheat seedlings during different time points. Pictures of 9-, 12- and 15-day-old winter wheat seedlings treated with 0 mg (control) and 25 mg PSI-362 in 75ml 1/10 MS agar medium for 1, 3, 6, and 9 days, respectively.

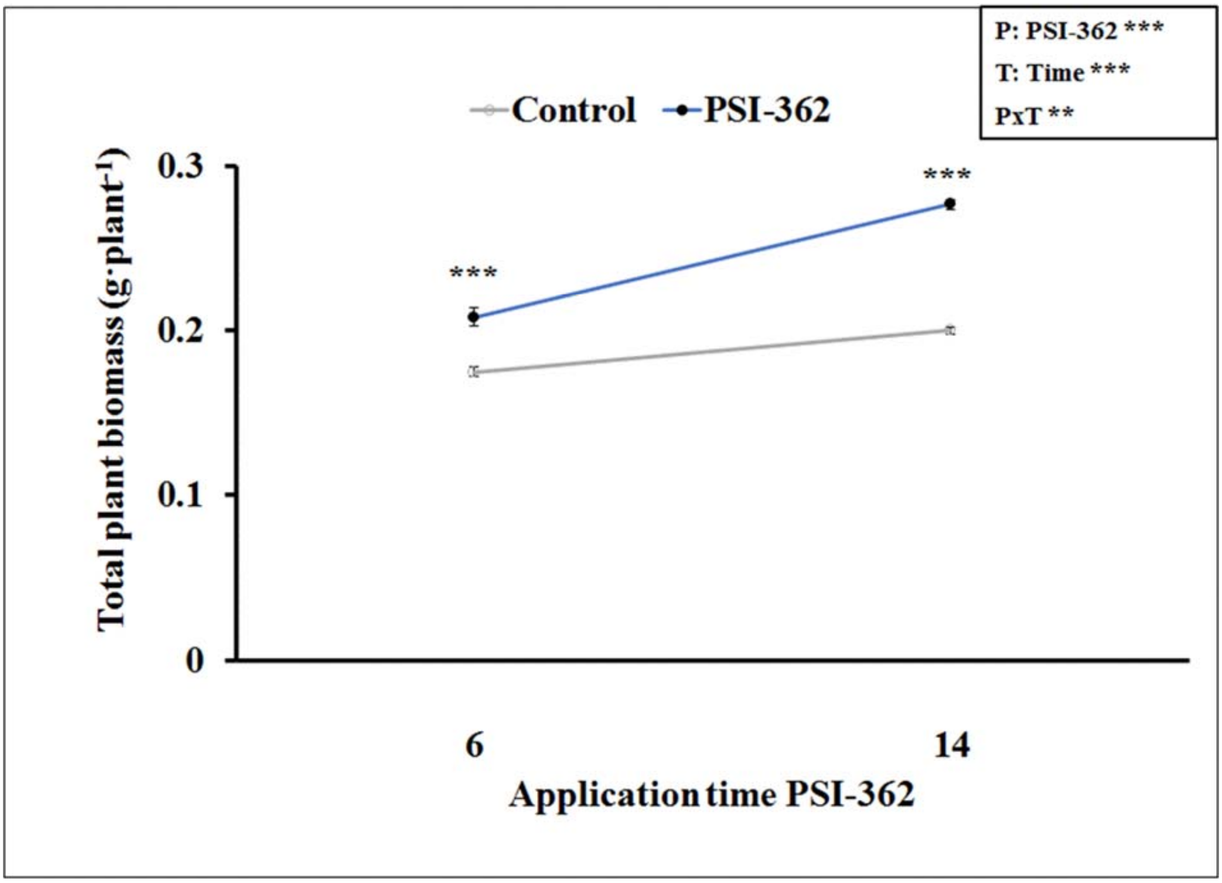


**Fig. S2.** PSI-362 increases plant biomass in winter wheat seedlings grown in nutrient starving conditions. 6-day-old winter wheat seedlings were treated with 0 mg (control) and 25 mg PSI-362 in 75 ml 0 MS agar medium for 6 and 14 days. Chart represents total plant biomass. Each treatment was performed using 9 independent biological replicates, with 8 plants per replicate. Since interaction PxT was significant (** $p \leq 0.01$), data were subjected to t-test, comparing PSI-362 treatment versus control within the same application time. In this case, means followed by asterisk (*** significant at $p \leq 0.001$) indicate statistically significant differences between control and PSI-362 treatment within the same application time.

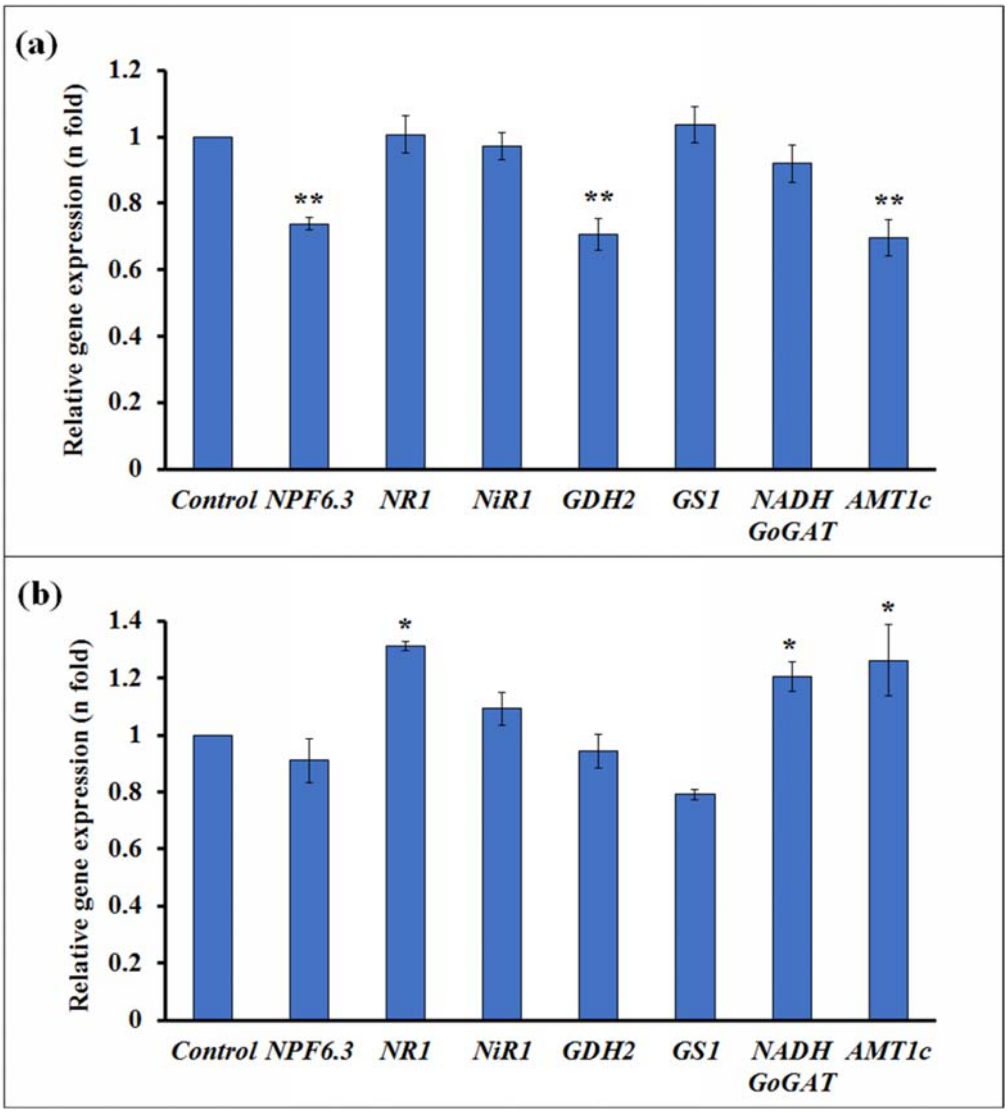


**Fig. S3.** Changes in relative gene expression in whole wheat seedlings with low PSI-362 concentration rate or using high N growth medium. 6-day-old winter wheat seedlings were treated with 0 mg (control) and 5 mg PSI-362 in 150 ml 1/10 MS agar medium for 6 days (a) or were treated with 0 mg (control) and 50 mg PSI-362 in 150 ml 1 MS agar medium for 6 days (b). Results are displayed as the relative log2 fold-change with respect to the TaEF1a gene expression levels. RNA extractions followed by RT-qPCR experiments were performed in at least 3 independent biological replicates. Means followed by asterisk indicate statistically significant differences between control and PSI-362 treatment within the same application time based on one-way ANOVA Student-Newman-Keuls' test (** and * significant at $p \leq 0.01$ and $p \leq 0.05$, respectively).

**Table S1.** Primers used for RT-qPCR analysis in winter wheat (cv. Graham)

| Primer | Sequence (5'-3') | Reference/ Accession |
|---|---|---|
| ***TaNPF6.3/AtNRT1.1 FW*** | CACGGGAGCAACGACGGCTG | (Buchner *et al.*, 2015) |
| ***TaNPF6.3/AtNRT1.1 REV*** | ATGCGTTTCTCCTTGTACACGTAG | (Buchner *et al.*, 2015) |
| ***TaNR1 FW*** | GGCCAATTCCTTCATCTCCTTCTG | (Buchner *et al.*, 2015) |
| ***TaNR1 REV*** | TACATGCACAGATTGATGCGTCGA | (Buchner *et al.*, 2015) |
| ***TaNIR FW*** | AACGTCAGGAACGACAAGGT | FJ527909.1 |
| ***TaNIR REV*** | GCGCCTTGGTCTCGATGAT | FJ527909.1 |
| ***TaGDH2 FW*** | AGGATGGGAGCATTCACCTTGG | (Buchner *et al.*, 2015) |
| ***TaGDH2 REV*** | GGATATAAGAACTCTCATCCACCACG | (Buchner *et al.*, 2015) |
| ***TaGS1 FW*** | ATGATCGCCGAGACCACCATCC | (Buchner *et al.*, 2015) |
| ***TaGS1 REV*** | TCGTCCAAATCCTCCACTGGCC | (Buchner *et al.*, 2015) |
| ***TaNADH-GoGAT FW*** | GCGGAAGTGCCATACCAATAC | (Li *et al.*, 2020) |
| ***TaNADH-GoGAT REV*** | AAAGTTAATGACATGCTCTGGTTCTC | (Li *et al.*, 2020) |
| ***TaAMT1_1c_FW*** | TTCTTGGCGCTCAAGAAG | (Li *et al.*, 2017) |
| ***TaAMT1_1c_REV*** | GCCGACCTGAGCATGAA | (Li *et al.*, 2017) |
| ***TaEF1a_FW*** | TGGTGTCATCAAGCCTGGTATGGT | (Li *et al.*, 2017) |
| ***TaEF1a_REV*** | ACTCATGGTGCATCTCAACGGACT | (Li *et al.*, 2017) |

**Table S2.** Effect of PSI-362 concentration applied for 6 days on NUE in winter wheat seedlings (cv. Graham) grown on reduced N media (1/10 MS).

| Parameter[1] | Treatment | | | |
|---|---|---|---|---|
| | Control | PSI-362 (5 mg) | PSI-362 (10 mg) | PSI-362 (50 mg) |
| **NUE (g FW·g$^{-1}$ N fertiliser)** | 130.95 ± 4.58 a | 144.84 ± 6.31 ab | 154.13 ± 0.73 b | 217.38 ± 3.02 c |

[1] Data are the means ± SE. Number of biological replicates (*n* = 9).

Different letters indicate statistical differences between treatments for $p \leq 0.05$ based on one-way ANOVA Student-Newman-Keuls' method

**Table S3.** Effect of PSI-362 applied for 6 days on NUE in winter wheat seedlings (cv. Graham) grown on reduced (1/10 MS) and saturated N media (1 MS).

| Parameter[1] | 1/10 MS | | 1 MS | |
|---|---|---|---|---|
| | Control | PSI-362 (50 mg) | Control | PSI-362 (50 mg) |
| **NUE (g FW·g$^{-1}$ N fertiliser)** | 123.02 ± 1.12 | 201.59 ± 3.74 * | 15.00 ± 0.22 | 19.48 ± 0.21 * |

[1] Data are the means ± SE. Number of biological replicates (*n* = 9).

Means followed by asterisk indicate statistically significant differences between control and PSI-362 treatment within the same N level based on t-test (*$p \leq 0.05$).

**Table S4.** Effect of PSI-362 applied during different time points on NUE in winter wheat seedlings (cv. Graham) grown on reduced N media (1/10 MS).

| Application time (days) | NUE (g FW·$g^{-1}$ N fertiliser)[1] | |
|---|---|---|
| | **Control** | **PSI-362 (50 mg)** |
| **1** | 115.87 ± 1.98 | 111.64 ± 2.41 |
| **3** | 195.63 ± 1.81 | 238.10 ± 10.39 * |
| **6** | 244.97 ± 2.63 | 353.44 ± 9.30 * |
| **9** | 279.89 ± 5.31 | 389.95 ± 7.50 * |

[1] Data are the means ± SE. Number of biological replicates ($n$ = 9).

Means followed by asterisk indicate statistically significant differences between control and PSI-362 treatment within the same application time based on t-test (*$p \leq 0.05$).

**Table S5.** Analysis of variance and mean comparisons for biochemical parameters in winter wheat seedlings (cv. Graham) grown on reduced N media (1/10 MS) and treated with PSI-362 during different time points.

| **Source of variance** | **$NH_4^+$ (µg·g$^{-1}$ FW)** | **FAA (mg·g$^{-1}$ FW)** | **Glu (mg·g$^{-1}$ FW)** | **Gln (mg·g$^{-1}$ FW)** | **Soluble prot. (mg·g$^{-1}$ FW)** | **Chl (a+b) (mg·g$^{-1}$ FW)** | **Carotenoids (mg·g$^{-1}$ FW)** |
|---|---|---|---|---|---|---|---|
| **PSI-362 (P)** | ** | ns | *** | ns | ** | ** | ** |
| **Time (T)** | *** | * | *** | *** | *** | *** | *** |
| **P x T** | * | ns | ** | ** | * | * | ns |
| **<u>PSI-362</u>** | | | | | | | |
| **Control** | 29.02 ± 2.10 b | 3.84 ± 0.10 a | 0.30 ± 0.01 a | 0.53 ± 0.08 a | 14.33 ± 0.58 a | 0.30 ± 0.02 a | 0.07 ± 0.01 a |
| **Treated** | 24.07 ± 3.45 a | 4.00 ± 0.09 a | 0.36 ± 0.01 b | 0.52 ± 0.11 a | 15.34 ± 0.86 b | 0.33 ± 0.02 b | 0.08 ± 0.01 b |
| **<u>Time</u>** | | | | | | | |
| **1 h** | 27.98 ± 2.56 b | 4.21 ± 0.09 b | 0.30 ± 0.01 a | 0.35 ± 0.01 a | 11.94 ± 0.31 a | 0.23 ± 0.01 a | 0.04 ± 0.00 a |
| **1 d** | 35.29 ± 0.50 c | 3.83 ± 0.08 a | 0.36 ± 0.02 c | 0.91 ± 0.03 b | 16.44 ± 0.33 b | 0.34 ± 0.01 b | 0.08 ± 0.00 b |
| **6 d** | 16.36 ± 2.44 a | 3.72 ± 0.07 a | 0.33 ± 0.01 b | 0.31 ± 0.02 a | 16.12 ± 0.47 b | 0.36 ± 0.01 b | 0.09 ± 0.00 b |
| **<u>P x T</u>** | | | | | | | |
| **Control + 1 h** | 28.49 ± 2.56 a | 4.22 ± 0.05 a | 0.30 ± 0.01 a | 0.37 ± 0.02 a | 12.10 ± 0.57 a | 0.24 ± 0.01 a | 0.04 ± 0.00 a |
| **Treated + 1 h** | 27.47 ± 2.41 a | 4.20 ± 0.18 a | 0.31 ± 0.01 a | 0.33 ± 0.01 a | 11.77 ± 0.19 a | 0.23 ± 0.01 a | 0.04 ± 0.00 b |
| **Control + 1 d** | 36.27 ± 0.52 a | 3.67 ± 0.07 a | 0.30 ± 0.00 a | 0.86 ± 0.01 a | 15.80 ± 0.34 a | 0.32 ± 0.01 a | 0.07 ± 0.00 a |
| **Treated + 1 d** | 34.32 ± 0.31 a | 4.00 ± 0.05 a | 0.41 ± 0.01 b | 0.96 ± 0.03 b | 17.08 ± 0.23 b | 0.37 ± 0.01 b | 0.09 ± 0.01 b |
| **Control + 6 d** | 22.30 ± 0.37 b | 3.64 ± 0.08 a | 0.31 ± 0.01 a | 0.35 ± 0.01 b | 15.08 ± 0.06 a | 0.34 ± 0.01 a | 0.08 ± 0.01 a |
| **Treated + 6 d** | 10.43 ± 0.38 a | 3.81 ± 0.09 a | 0.36 ± 0.00 b | 0.27 ± 0.02 a | 17.15 ± 0.41 b | 0.39 ± 0.01 b | 0.10 ± 0.01 b |

ns, *, **, *** Non-significant or significant at $p \leq 0.05$, $p \leq 0.01$, and $p \leq 0.001$, respectively. Different letters indicate statistical differences for $p \leq 0.05$ based on t-test (P and P x T) or Student–Newman–Keuls' method (T). Data are the means ± SE. Number of biological replicates (n ≥ 4).

**Table S6.** Analysis of variance and mean comparisons for biochemical parameters in winter wheat seedlings (cv. Graham) treated with PSI-362 for 6 days and growing under two different N rates.

| Source of variance | $NH_4^+$ (µg·g$^{-1}$ FW) | FAA (mg·g$^{-1}$ FW) | Glu (mg·g$^{-1}$ FW) | Gln (mg·g$^{-1}$ FW) | Sol. prot. (mg·g$^{-1}$ FW) | Chl (a+b) (mg·g$^{-1}$ FW) | Carot. (mg·g$^{-1}$ FW) |
|---|---|---|---|---|---|---|---|
| **PSI-362 (P)** | ** | ns | ns | ** | * | ns | ns |
| **N rate (N)** | *** | ** | *** | *** | *** | *** | * |
| **P x N** | *** | ns | ** | ns | ns | * | ns |
| **PSI-362** | | | | | | | |
| **Control** | 57.68 ± 11.58 a | 4.60 ± 0.15 a | 0.43 ± 0.05 a | 0.86 ± 0.18 b | 17.11 ± 0.78 a | 0.43 ± 0.02 a | 0.10 ± 0.00 a |
| **Treated** | 61.89 ± 20.25 b | 4.36 ± 0.26 a | 0.44 ± 0.02 a | 0.76 ± 0.17 a | 18.43 ± 0.47 b | 0.44 ± 0.01 a | 0.10 ± 0.01 a |
| **N rate** | | | | | | | |
| **1/10x** | 20.82 ± 3.50 a | 4.89 ± 0.14 b | 0.36 ± 0.03 a | 0.39 ± 0.02 a | 16.42 ± 0.50 a | 0.40 ± 0.01 a | 0.09 ± 0.00 a |
| **1x** | 98.75 ± 5.23 b | 4.07 ± 0.14 a | 0.51 ± 0.03 b | 1.22 ± 0.04 b | 19.13 ± 0.33 b | 0.48 ± 0.01 b | 0.11 ± 0.00 b |
| **P x N** | | | | | | | |
| **Control + 1/10x** | 29.34 ± 0.63 b | 4.87 ± 0.17 a | 0.30 ± 0.00 a | 0.42 ± 0.01 a | 15.40 ± 0.30 a | 0.38 ± 0.01 a | 0.09 ± 0.00 a |
| **Treated + 1/10x** | 12.30 ± 0.22 a | 4.91 ± 0.21 a | 0.42 ± 0.03 b | 0.37 ± 0.02 a | 17.44 ± 0.46 b | 0.42 ± 0.01 b | 0.09 ± 0.01 a |
| **Control + 1x** | 86.02 ± 0.87 a | 4.33 ± 0.09 a | 0.55 ± 0.03 b | 1.30 ± 0.03 b | 18.82 ± 0.60 a | 0.49 ± 0.00 a | 0.11 ± 0.00 a |
| **Treated + 1x** | 111.47 ± 0.93 b | 3.82 ± 0.17 a | 0.46 ± 0.01 a | 1.15 ± 0.01 a | 19.43 ± 0.13 a | 0.46 ± 0.02 a | 0.11 ± 0.01 a |

ns, *, **, *** Non-significant or significant at $p \leq 0.05$, $p \leq 0.01$, and $p \leq 0.001$, respectively. Different letters indicate statistical differences for $p \leq 0.05$ based on t-test (P, N, and P x N). Data are the means ± SE. Number of biological replicates (n ≥ 4).

### References for supporting information

**Buchner P, Tausz M, Ford R, Leo A, Fitzgerald GJ, Hawkesford MJ, Tausz-Posch S. 2015.** Expression patterns of C- and N-metabolism related genes in wheat are changed during senescence under elevated CO2 in dry-land agriculture. *Plant Sci* **236**: 239-249.

**Li T, Liao K, Xu X, Gao Y, Wang Z, Zhu X, Jia B, Xuan Y. 2017.** Wheat Ammonium Transporter (AMT) Gene Family: Diversity and Possible Role in Host–Pathogen Interaction with Stem Rust. *Frontiers in plant science* **8**(1637).

**Li W, He X, Chen Y, Jing Y, Shen C, Yang J, Teng W, Zhao X, Hu W, Hu M, et al. 2020.** A wheat transcription factor positively sets seed vigour by regulating the grain nitrate signal. *New Phytologist* **225**(4): 1667-1680.
